# Inosine misincorporation into mRNA triggers the integrated stress response and activates an innate immune gene expression signature

**DOI:** 10.64898/2026.01.16.699979

**Authors:** Jacob H. Schroader, Naa N. Adade, Emmanuel E. Adade, Mark A. Bratslavsky, Hannah K. Shorrock, Bradley R. Smith, Ting Zhou, Steven Lotz, Taylor Bertucci, Sally Temple, Alex M. Valm, Cara T. Pager, J. Andrew Berglund, Mark T. Handley, Gabriele Fuchs, Kaalak Reddy

## Abstract

The inosine triphosphate pyrophosphatase (ITPase) enzyme restricts levels of the non-canonical nucleotides (deoxy)inosine triphosphate (dITP/ITP) and prevents their aberrant misincorporation into nucleic acids. ITPase deficiency is associated with dilated cardiomyopathy and epileptic encephalopathy in humans and is usually fatal in infancy. It leads to pronounced inosine misincorporation into RNA but the cellular consequences of this misincorporation are not well understood and the pathogenic basis of ITPase deficiency remains unknown. Here we show that cellular transfection of mRNA with inosine misincorporation activates the integrated stress response (ISR) with an innate immune gene expression signature. This stress response triggers stress granule formation and is modulated by the double stranded RNA sensor protein Kinase R (PKR). Inosine nucleoside treatment of ITPase-deficient cells leads to endogenous inosine misincorporation into mRNA and activation of the ISR. Further, differentiation of human ITPase-deficient induced pluripotent stem cells into neurons results in a low-level stress response. Thus, our study establishes inosine misincorporation into mRNA as an unappreciated form of cellular stress. This is normally prevented by the ITPase enzyme, with implications for the pathogenesis of ITPase deficiency.

## INTRODUCTION

Proper cellular function requires stringent maintenance of nucleotide pools, in terms of their levels as well as their composition^1,2^. Failure to prevent accumulation of damaged and non-canonical nucleotides can have severe biological consequences. The ITPase enzyme preserves the integrity of cellular nucleotide pools by hydrolyzing the non-canonical purine nucleotides dITP and ITP^3–6^. ITPase is necessary because inosine monophosphate (IMP) is an intermediate in the generation of adenosine and guanosine monophosphates (AMP and GMP, respectively) during purine nucleotide biosynthesis, and adventitious phosphorylation of (d)IMP can generate (d)ITP^3–6^. Deamination of (d)ATP may also generate (d)ITP^3–6^.

The importance of ITPase is underscored by the conservation of orthologous enzymes in all domains of life^3,5^. Loss of this enzyme in a range of model organisms including *E. coli, Saccharomyces cerevisiae*, *Arabidopsis thaliana* and mice results in distinct and overlapping biological outcomes^7–13^. We and others previously identified biallelic loss-of-function variants in the *ITPA* gene encoding ITPase, leading to absent or compromised enzyme activity, as the cause of a fatal multisystem disorder (MIM 616647) characterized by epileptic encephalopathy and dilated cardiomyopathy^14–19^. *Itpa*-null mouse models reveal concordant features including seizures and structural heart abnormalities^11,12^.

The tissues of *Itpa*-null mice reveal substantial inosine misincorporation into RNA, with an incorporation frequency of up to ∼ 1% of AMP or ∼ 1 in 400 nucleotides of total RNA in the heart ^11,12,15^. We did not detect deoxyinosine within genomic DNA of *Itpa*-null mouse cells or tissues, nor did they exhibit any evidence of nuclear or mitochondrial DNA damage or instability^15^. This is consistent with active DNA repair pathways functioning to recognize and repair dITP misincorporation to preserve genome integrity^20,21^. While inosine is a necessary RNA modification, its introduction by adenosine deaminases in A-to-I editing is a highly regulated process^22–25^. Furthermore, the presence of inosine within mRNA codons has context-dependent effects on translation, not always behaving as a guanosine mimic^26^. Thus, misincorporation of inosine, with it’s distinct hydrogen bonding potential^27,28^, in place of the canonical nucleotides within RNA, could have detrimental effects on multiple facets of the RNA lifecycle^3^, therefore necessitating an enzyme, ITPase, that restricts the accumulation of inosine nucleotides. While the biochemical function of ITPase enzyme is well understood, the pathogenic basis of ITPase deficiency and the cellular consequences of inosine misincorporation into RNA remain unknown.

We previously found that inosine misincorporation into mRNA hinders translation of the encoded protein *in vitro* and in cells^29^. However, the broader cellular consequences of widespread mRNA inosine misincorporation remain poorly understood. Here, we show that inosine misincorporation into mRNA triggers stress granule formation and activates the integrated stress response (ISR) with a gene expression signature that resembles antiviral innate immunity. Our data suggests that PKR is a sensor of inosine misincorporation into mRNA. In the absence of the ITPase enzyme, treatment of cells with inosine nucleoside results in misincorporation into mRNA and activation of the ISR. Therefore, there is a response to endogenously generated inosine-containing mRNA. Furthermore, we show that ITPase-deficient human induced pluripotent stem cells (iPSCs) exhibit gene expression indicative of a low-level stress response upon differentiation into neurons. Together, these findings establish the importance of preventing inosine misincorporation into mRNA and reveal a potential mechanism that could contribute to pathogenesis in ITPase deficiency.

## RESULTS

### mRNA containing stochastic inosine misincorporation triggers an innate immune gene expression signature in cells

To evaluate the global cellular response to mRNA with misincorporated inosine, we performed RNA sequencing of cells following transfection with control, and inosine-containing firefly luciferase (FLuc) mRNA. Wildtype H9c2 rat cardiomyoblast cells were used, in which we previously reported reduced translation of transfected FLuc mRNA due to inosine misincorporation^29^. T7 *in vitro* transcription was used to generate capped FLuc mRNA in the presence of 10 mM of each canonical NTP plus 0, 0.1 or 1 mM ITP (Figure 1A). This system enables us to quantitatively titrate inosine misincorporation into mRNA with previous mass spectrometry indicating that it leads to stochastic inosine misincorporation at a frequency of 0, 1 in ∼ 9000 and 1 in ∼ 1000 nucleotides, respectively^29^. These misincorporation rates are within the range previously reported in murine models of ITPase deficiency^11,12,15^. Previous Oxford Nanopore direct RNA sequencing revealed that inosine is stochastically misincorporated in place of any of the canonical nucleotides, albeit with a preference for G>C>A>U^29^. These FLuc RNA preparations, referred to as 0 (control), 0.1 and 1 mM ITP, were polyadenylated, column purified and then transfected into H9c2 cells (Figure 1A).

**Figure 1.**
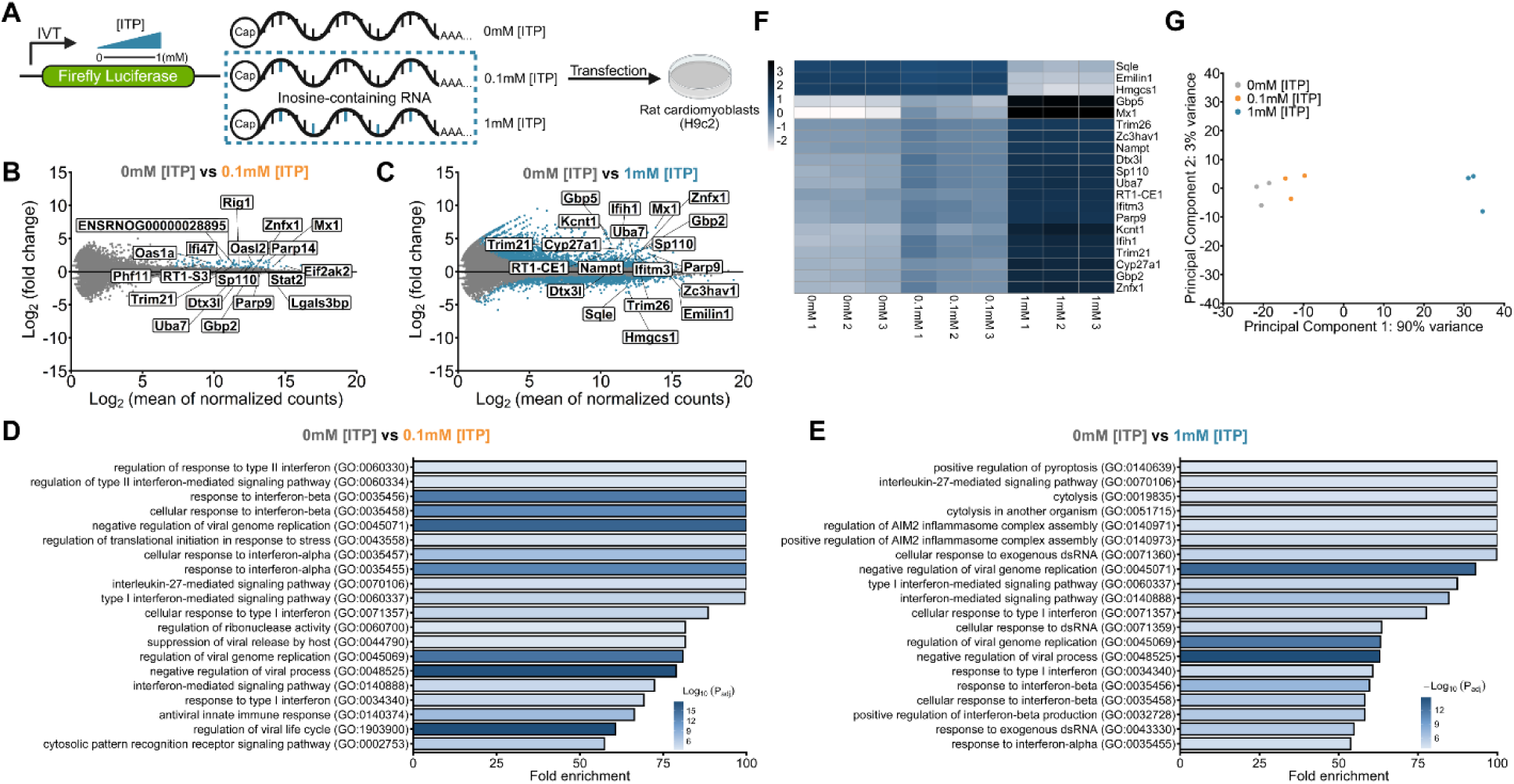
Transfection of H9c2 rat cardiomyoblast cells with mRNA containing inosine misincorporation leads to upregulation of innate immune gene expression. **(A)** Inosine containing mRNA is generated through *in vitro* transcription of linearized firefly luciferase (FLuc) template using T7 phage RNA polymerase in the presence of 10 mM concentration of each canonical nucleotide and 0, 0.1 and 1 mM ITP in the nucleotide pool leading to inosine misincorporation. Control and inosine mRNA was then transfected into cells. **(B)** Illumina RNA sequencing was performed on ribosomal-depleted cellular RNA 24 hours post transfection in triplicate. MA plots showing differentially expressed genes in 0 vs. 0.1 mM [ITP] mRNA transfections and **(C)** 0 vs. 1 mM [ITP] with the most significantly altered genes highlighted. p-adjusted values < 0.05. **(D)** Gene ontology analysis of the 20 most significant genes in the 0 vs. 0.1 mM [ITP] and **(E)** 0 vs 1 mM [ITP] comparisons. p-adjusted values < 0.05. **(F)** Heatmap of regularized log transformed expression of top 20 significant genes in the 0 vs. 1 mM [ITP] comparison for all libraries. **(G)** Unfiltered principal component analysis for all libraries.

We first confirmed a dose-dependent reduction in firefly luciferase activity due to impaired translation owing to inosine misincorporation within the mRNA^29^ (Figures S1A, S1B). Following transfection of cells with the control, 0.1 and 1 mM inosine mRNA, total cellular RNA was extracted, depleted of ribosomal RNA and subjected to Illumina RNA sequencing. Gene expression analysis using DESeq2^30^ revealed an unexpected and striking dose-dependent innate immune gene expression signature 24 hours following transfection with the inosine-containing mRNAs (Figures 1B, 1C). Gene ontology (GO) analysis identified significant enrichment of terms related to innate immunity, interferon signalling and response to virus from both 0.1 and 1mM inosine mRNA transfections (Figures 1C, 1D). Notably, the 1 mM inosine condition was enriched for terms related to cell death such as “pyroptosis” and “cytolysis” signifying a severe cellular response to mRNA containing inosine misincorporation (Figure 1E). Consistently, this was reflected in a significant effect on cellular viability (Figures S1A, S1C). Significantly altered genes in both 0.1 and 1 mM inosine groups were biased towards upregulation with a greater magnitude of change in the 1 mM group (Figure 1F). Upregulated genes included well-known viral sensing and interferon-stimulated genes (ISGs) linked to innate immunity, such as *Gbp5*, *Mx1* and *Ifih1* ^31–33^. Furthermore, unfiltered principal component analysis clustered libraries according to treatment group, with the 0.1 mM inosine group intermediate to control and 1 mM inosine groups (Figure 1G). This highlights the discrete and graded transcriptomic response to inosine misincorporation within mRNA (Figure 1G).

### mRNA with inosine misincorporation activates the integrated stress response

From the RNAseq dataset, we selected a panel of significantly upregulated genes that encode factors linked to innate immunity^33^, comprised of *Adar*, *Eif2ak2* encoding PKR, *Ifih1* encoding MDA5 and *Rig1* (Figure S2). We transfected H9c2 cells with control or 1mM inosine FLuc mRNA and monitored expression of the panel over a 72-hour time course (Figure 2A). Confirming RNAseq findings, there was significant upregulation of each factor at 24 hours. However, expression diminished to basal levels by 72 hours. Higher levels of inosine FLuc mRNA than the control mRNA were maintained in cells over the time course (Figure 2B), whereas FLuc protein activity in inosine FLuc mRNA-transfected cells was orders of magnitude lower (Figure 2C).

**Figure 2.**
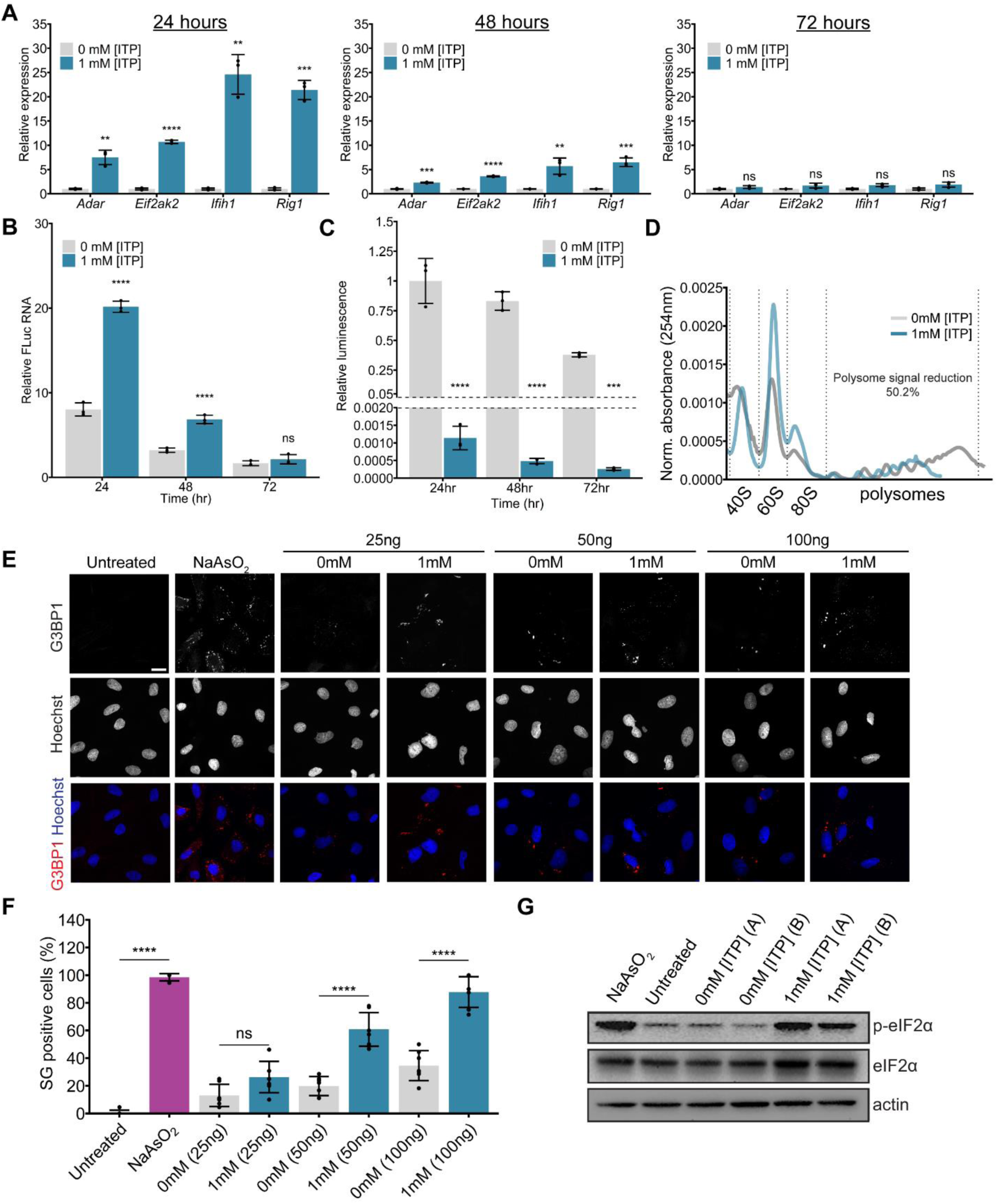
Induction of stress granule formation and activation of the integrated stress response upon transfection of mRNA with inosine misincorporation. **(A)** Expression of *Adar, Eif2ak2*, *Ifih1* & *Rig1* transcripts relative to *Gapdh* following transfection of 0 and 1 mM [ITP] FLuc mRNA over a 72 hour time-course and normalized to the 0 mM [ITP] control mRNA. **(B)** FLuc RNA levels relative to *Gapdh* over a 72 hour time-course. **(C)** Relative luminescence normalized to 0 mM [ITP] control FLuc RNA. Mean +/- SD, n = 3 experimental replicates. **(D)** Polysome profiles (normalized AUC absorbance) with calculated polysome signal reduction values ([treatment polysome signal-control polysome signal]/control polysome signal X 100). Following signal reduction quantification, x-axis dilation was used on 1mM [ITP] traces to align 80S peaks. **(E)** Immunofluorescence for G3BP1 and Hoechst nuclei staining following transfection of 25, 50 or 100 ng of either 0 or 1mM [ITP] FLuc mRNA for 24 hours. Scale bar = 10 μm. **(F)** Quantification of proportion of stress granule (SG) positive cells. Mean +/- SD, n = 7 fields of view with 49-165 cells per group. **(G)** Immunoblot of (phospho) p-eIF2α, eIF2α and actin as loading control following transfection of 0 and 1 mM [ITP] FLuc mRNA for 24 hours. A and B represent independent experimental replicates per group. One-way ANOVA/Dunnett’s multiple comparisons test, **P < 0.01, ***P < 0.001, ****P < 0.0001, ns – not significant.

To ensure that the transcriptional response to inosine-containing mRNA was not dependent on a particular mRNA sequence, we repeated our experiments using *Renilla* luciferase (RLuc) mRNA and observed similar results (Figure S3). Likewise, to ensure that the effects were not restricted to H9c2 cells, control and inosine FLuc mRNA were transfected into A549 human lung carcinoma cells with similar results (Figures S4A-C). Taken together, mRNA containing misincorporated inosine triggers a cellular innate immune response.

Given the transcriptomic response and enrichment for terms related to antiviral innate immunity, we wondered if there was a global block to translation, which is observed with viral infection. To assess translation, we performed polysome profiling after transfecting H9c2 cells with either control or 1mM inosine FLuc mRNA (Figure 2D). Area under the curve quantification showed that the proportion of polysomes (actively translating ribosomes) was 50% lower in the cells transfected with the 1 mM inosine FLuc mRNA (Figure 2D). This indicates that global translation is reduced in cells transfected with inosine-containing mRNA.

The greater abundance/reduced turn-over of inosine-containing mRNA compared to control mRNA (Figure 2B) raised the possibility that it might be differentially stabilized through stress granule formation. Stress granule formation would also be consistent with the reduced functional translation of inosine-containing FLuc mRNA (Figure 2C), and the observed reduction of translation globally (Figure 2D). To test our hypothesis, we carried out immunofluorescence and confocal microscopy for the stress granule marker G3BP1^34^. Stress granules were essentially absent in untreated cells but present in virtually all cells upon treatment with sodium arsenite as a positive control (Figure 2E). Upon mRNA transfections, there was a clear and significant increase in the percentage of stress granule positive cells following transfection with inosine containing mRNA, compared to control mRNA, at a series of different mRNA concentrations (Figure 2F). Stress granule formation was induced with inosine mRNAs at comparable concentrations to those producing the transcriptional response and translational repression.

Stress granule formation and global translation blockade is associated with activation of the integrated stress response (ISR)^34^. Moreover, analysis of the RNAseq data showed a significant dose-dependent upregulation of *Atf4,* a key regulator of the ISR, together with several of its downstream targets including *Atf3, Ddit3,* and *Trib3*^35,36^, following transfection of H9c2 cells with 0.1 and 1 mM inosine mRNA (Figure S5). We therefore evaluated eIF2α phosphorylation, the hallmark signal of the ISR, using immunoblotting (Figure 2G). There was a strongly elevated phospho-eIF2α signal following inosine mRNA transfection, comparable to that produced by treatment of the cells with (1 mM) sodium arsenite. This was not evident in either untreated cells or cells transfected with control mRNA (Figure 2G). Taken together, our findings indicate that mRNA containing misincorporated inosine triggers stress granule formation, activates the integrated stress response, and is associated with an innate immune gene expression signature.

### PKR modulates the cellular stress response to mRNA with misincorporated inosine

The four kinases that are known to phosphorylate eIF2α leading to the ISR are GCN2, PKR, HRI and PERK^35^. As our RNAseq and subsequent qPCR had shown consistent upregulation of *Eif2ak2* - encoding PKR – with inosine mRNA transfections, we hypothesized that PKR was responsible for eIF2α phosphorylation produced by these transfections. We were unable to directly confirm phospho-PKR levels in the H9c2 cells owing to the limitation of phospho-PKR antibodies for murine samples^37^. Therefore, we evaluated the role of PKR using imidazolo-oxindole PKR Inhibitor C16 (Sigma 527450). We measured expression of our panel of stress response factors, together with cellular viability in H9c2 cells transfected with control or 1 mM inosine FLuc mRNA, with and without PKR inhibition (Figure 3A). Inosine mRNA-mediated upregulation of *Adar1, Atf4, Eif2ak2, Ifih1* and *Rig1*, were significantly attenuated with the PKR inhibitor which was not the case in control mRNA transfections (Figure 3B). PKR inhibitor also significantly rescued the reduced cellular viability caused by the inosine mRNA (Figure 3C). We directly confirmed PKR activation in response to inosine mRNA in the human A549 cell line using a phospho-PKR antibody (Figure S4D). There was a clear increase in phospho-PKR signal following transfection with inosine mRNA compared to control mRNA in these cells. These findings implicate PKR as a potential mediator of the cellular stress response to inosine misincorporation into mRNA.

**Figure 3.**
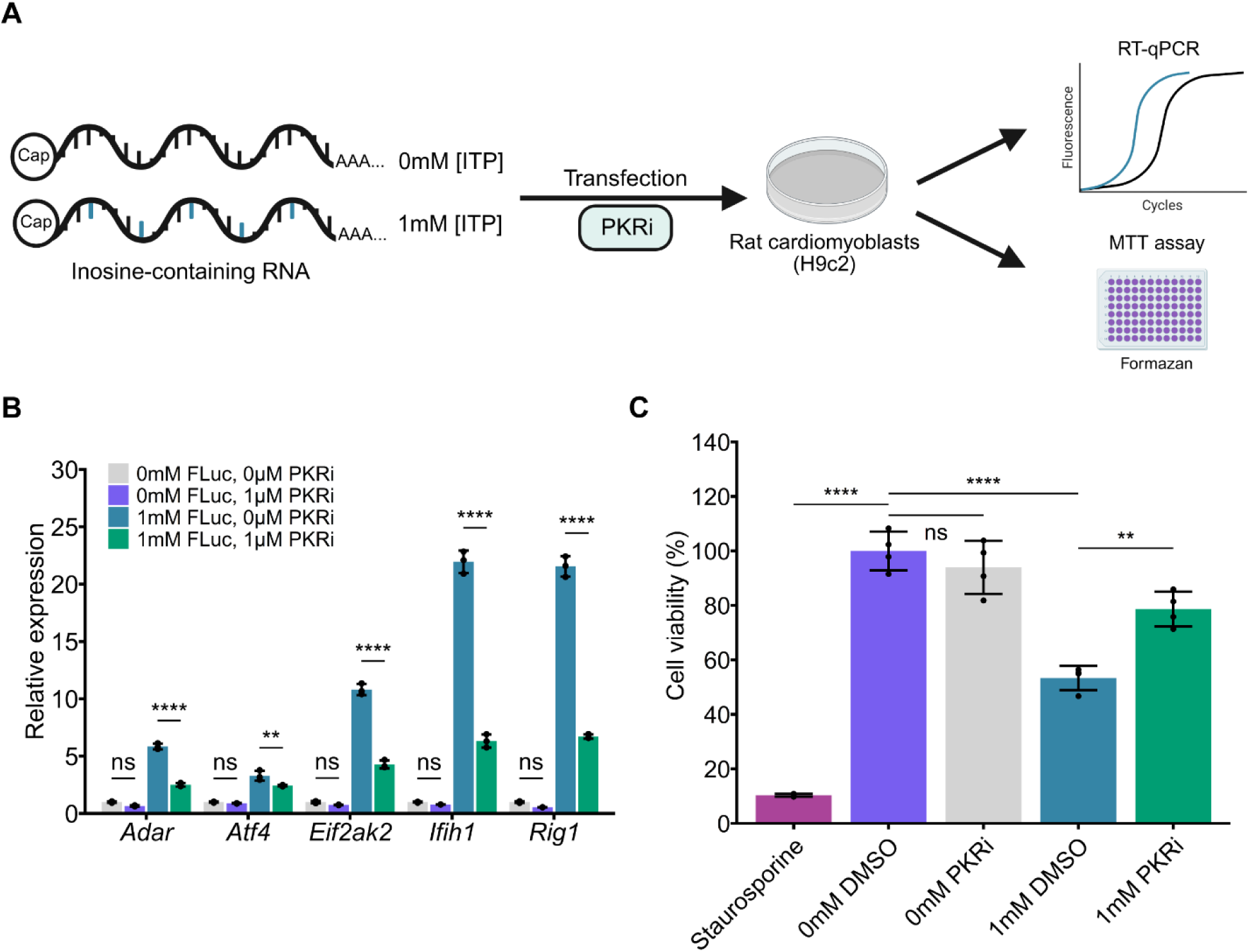
PKR modulates the cellular response to inosine mRNA. **(A)** Treatment of H9c2 cells with PKR inhibitor C16 (PKRi), 6 hours post-transfection of 0 or 1 mM [ITP] FLuc mRNA totaling 30 hours followed by luciferase assay, RT-qPCR, and cell viability. **(B)** Relative expression of *Adar*, *Atf4, Ifih1*, *Eif2ak2* and *Rig1*,. Mean +/- SD, n = 3 experimental replicates, Student’s t-test with Holm-Bonferroni correction, **P < 0.01, ****P < 0.0001, ns – not significant. **(C)** Cell viability represented as a percent relative to the untreated (DMSO) group. Mean +/- SD, n = 4 experimental replicates, One-way ANOVA/Tukey’s HSD, **P < 0.01, ****P < 0.0001, ns – not significant.

### mRNA determinants of the stress response to inosine misincorporation

Because PKR is activated by direct binding to dsRNA, we tested whether inosine RNA preparations independent of an mRNA context would induce the cellular stress response. We transfected control and 1 mM inosine FLuc and RLuc RNA without a cap or polyA tail into H9c2 cells and evaluated expression of our candidate stress response gene panel. There was no response to these RNAs (Figure S6). These data suggest that any potential dsRNA structures that may be introduced via inosine misincorporation were not sufficient to elicit the stress response in the absence of an mRNA context.

Given the cellular response to inosine misincorporation within a capped and polyadenylated mRNA, we next tested if disrupting translation by altering the start codon of the mRNA would have any effect. We generated a set of FLuc start codon mutant plasmids with a near-cognate (CTG), a non-cognate (AGG) and a stop codon (TGA) in place of the canonical ATG start. These constructs were used to generate capped and polyadenylated control and 1 mM inosine FLuc mRNA, which were then transfected into H9c2 cells (Figure 4A). We assessed effects on translation by measuring luminescence, revealing a clear reduction in both control and inosine mRNA with any of the start codon mutants, as expected, and with a greater reduction evident in the inosine mRNAs compared to the controls, consistent with our observations for the canonical AUG mRNA (Figure 4B). Strikingly, in contrast to their effects on translation, the pattern of upregulation of the stress response gene panel was unaffected by the start codon mutants (Figure 4C). Inosine-containing mRNA produced the characteristic response in each case, but it was absent with the control mRNAs (Figure 4C).

**Figure 4.**
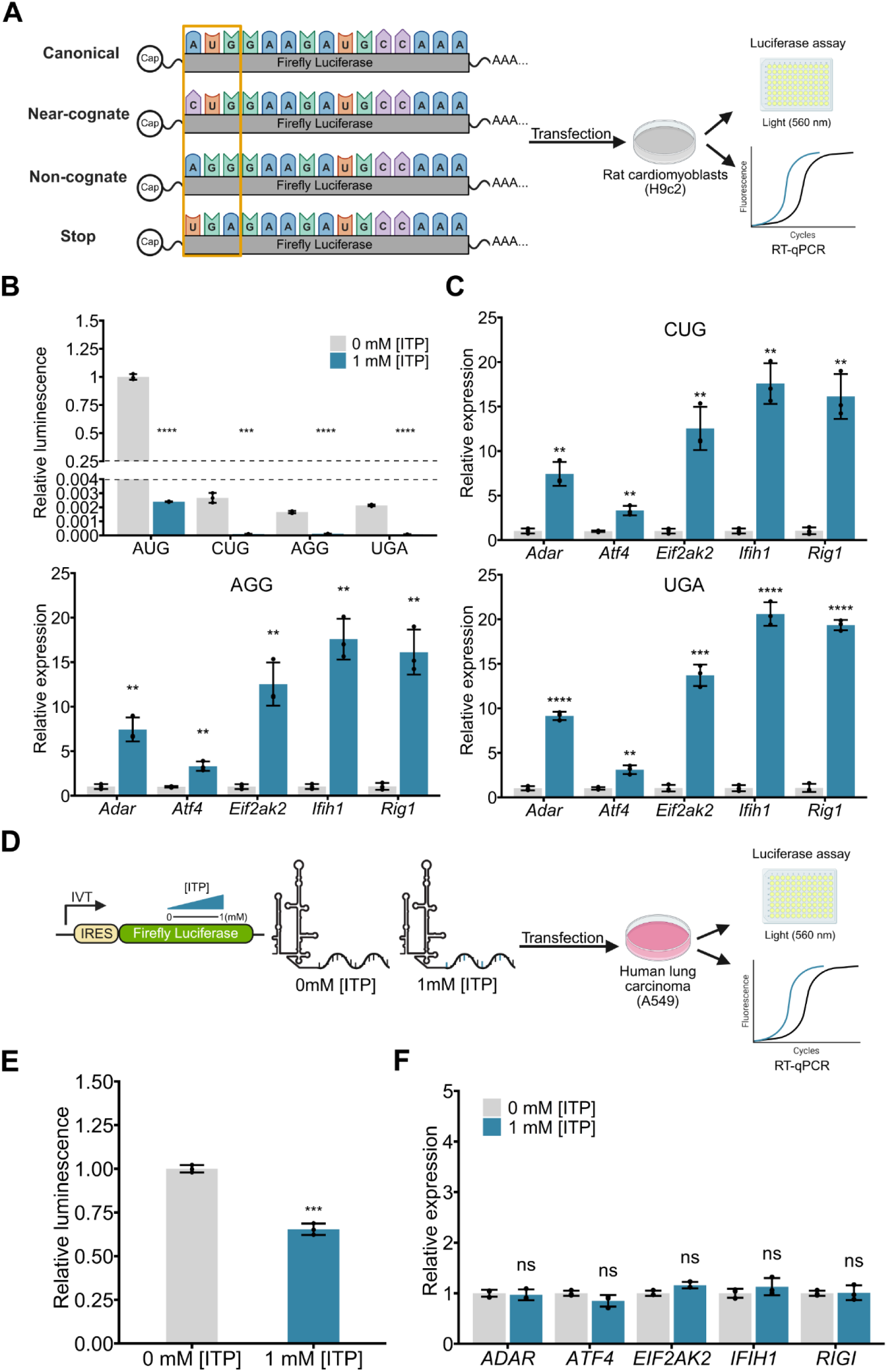
Disrupting translation of inosine-containing RNA does not influence the cellular stress response. **(A)** Near-cognate, non-cognate, and stop codon versions of 0 or 1 mM [ITP] FLuc mRNA were transfected into H9c2 cells for 30 hours and evaluated for luminescence and expresison of our stress response gene panel. **(B)** Luminescence relative to AUG control FLuc mRNA (0 mM). Mean +/- SD, n = 3 experimental replicates, Student’s t-test with Holm-Bonferroni correction, ***P < 0.001, ****P < 0.0001. **(C)** Expression of *Adar, Atf4, Eif2ak2*, *Ifih1* & *Rig1* transcripts relative to *Gapdh* for each start codon mutation (CUG, AGG, UGA). Mean +/-SD, n = 3, Student’s t-test with Holm-Bonferroni correction for each factor, *P < 0.05, **P < 0.01, ***P < 0.001, ****P < 0.0001. **(D)** Transfection of 0 or 1 mM [ITP] HCV-IRES FLuc RNA into A549 human lung carcinoma cells for 18 hours followed by luciferase assay and RT-qPCR. **(E)** Luminescence relative to control FLuc RNA (0 mM). Mean +/- SD, n = 3 experimental replicates, Student’s t-test, ***P < 0.001. **(F)** Expression of *ADAR*, *ATF4, EIF2AK2*, *IFIH1* & *RIGI* transcripts relative to *GAPDH*. Mean +/- SD, n = 3 experimental replicates, Student’s t-test with Holm-Bonferroni correction, ns – not significant.

mRNA capping and polyadenylation are associated with canonical translation initiation. We therefore wondered if engineering inosine-containing mRNA with the capacity for non-canonical translation-initiation would enable it to induce a similar stress response. To this end, we tested the effects of an internal ribosome entry site (IRES) RNA substrate which utilizes a cap-independent non-canonical form of translation as observed with certain RNA viruses^38^. We fused an upstream hepatitis C virus (HCV) IRES sequence to FLuc sequence lacking a canonical start codon. This is sufficient to initiate IRES-meditated translation in cells^39–41^. As previously, RNA was transcribed by T7 *in vitro* transcription with no (0mM) or 1 mM ITP in the reaction. Transfections were performed with A549 human lung carcinoma cells, which are IRES translation competent, then luminescence and expression of our stress response gene panel were evaluated (Figure 4D). While there was a significant reduction in translation of functional FLuc from the inosine-containing HCV IRES RNA compared to the control mRNA, there was no change in expression of our gene panel with the inosine HCV IRES RNA (Figures 4E, 4F). Taken together, these results indicate that a discrete cellular stress response to inosine misincorporation results from an mRNA context but is not influenced by active translation.

### Inosine nucleoside treatment triggers the integrated stress response in ITPase-deficient cells

Having established that transfections of inosine-containing mRNA result in upregulation of innate immune gene expression and activation of the integrated stress response, we next addressed whether endogenous misincorporation of inosine nucleotide into cellular RNA evokes a similar response. To investigate this, we utilized ITPase-deficient H9c2 rat cardiomyoblasts that misincorporate inosine within endogenous RNA, together with isogenic wild-type controls in which this misincorporation is restricted^29^. Further, we treated these cells with inosine nucleoside in the culture media, which is taken up by the cells and phosphorylated into nucleotide form. Wild-type cells with functional ITPase enzyme restrict the accumulation of ITP and the subsequent misincorporation into RNA. In ITPase-deficient cells, we predicted elevated ITP generation and inosine misincorporation into RNA. To validate this prediction, wild-type and *Itpa*-null rat H9c2 cells were treated with inosine nucleoside at 0, 1 and 10 mM in the culture media for 24 hours and then total RNA was extracted, digested into nucleosides, and analyzed using mass spectrometry (LC-MS). Consistent with our prediction, there was no detectable accumulation of inosine within the bulk RNA from wild-type cells (Figure S7). However, in *Itpa*-null cells, we observed elevated inosine content, proportional to the concentration of inosine nucleoside in the cell culture media (Figure S7). There was an inosine incorporation frequency of ∼ 1 in 25 000, 1 in 20 000 and 1 in 13 000 bases within total RNA in the 0, 1, and 10 mM treatments, respectively, reflecting an approximately 2-fold increase in inosine misincorporation with 10 mM inosine nucleoside treatment of the *Itpa*-null cells (Figure S7).

To confirm inosine misincorporation into mRNA, we next performed Oxford Nanopore Technologies (ONT) PromethION direct RNA sequencing on RNA from wild-type and *Itpa*-null cells treated with 0 (control) or 10 mM inosine nucleoside in the culture media (Figure 5A). Although the ONT basecaller Dorado enables inosine basecalling from direct RNA sequencing, it is only configured for inosine detection at A positions within the context of A-to-I editing, thus missing inosine incorporation at other nucleotide positions. However, we previously found that inosine gets misincorporated stochastically and in place of any nucleotide during T7 RNA polymerase transcription *in vitro*^29^ using ONT direct RNA sequencing by measuring base substitution frequency as a proxy for inosine misincorporation^42^. Because inosine within mRNA generates a unique current during ONT direct RNA sequencing, those positions will be miscalled by the basecaller, leading to a mismatch relative to the reference genome (Figure 5A). We used this approach, predicting that *Itpa*-null cells would exhibit greater base substitution rates due to inosine misincorporation into mRNA.

**Figure 5.**
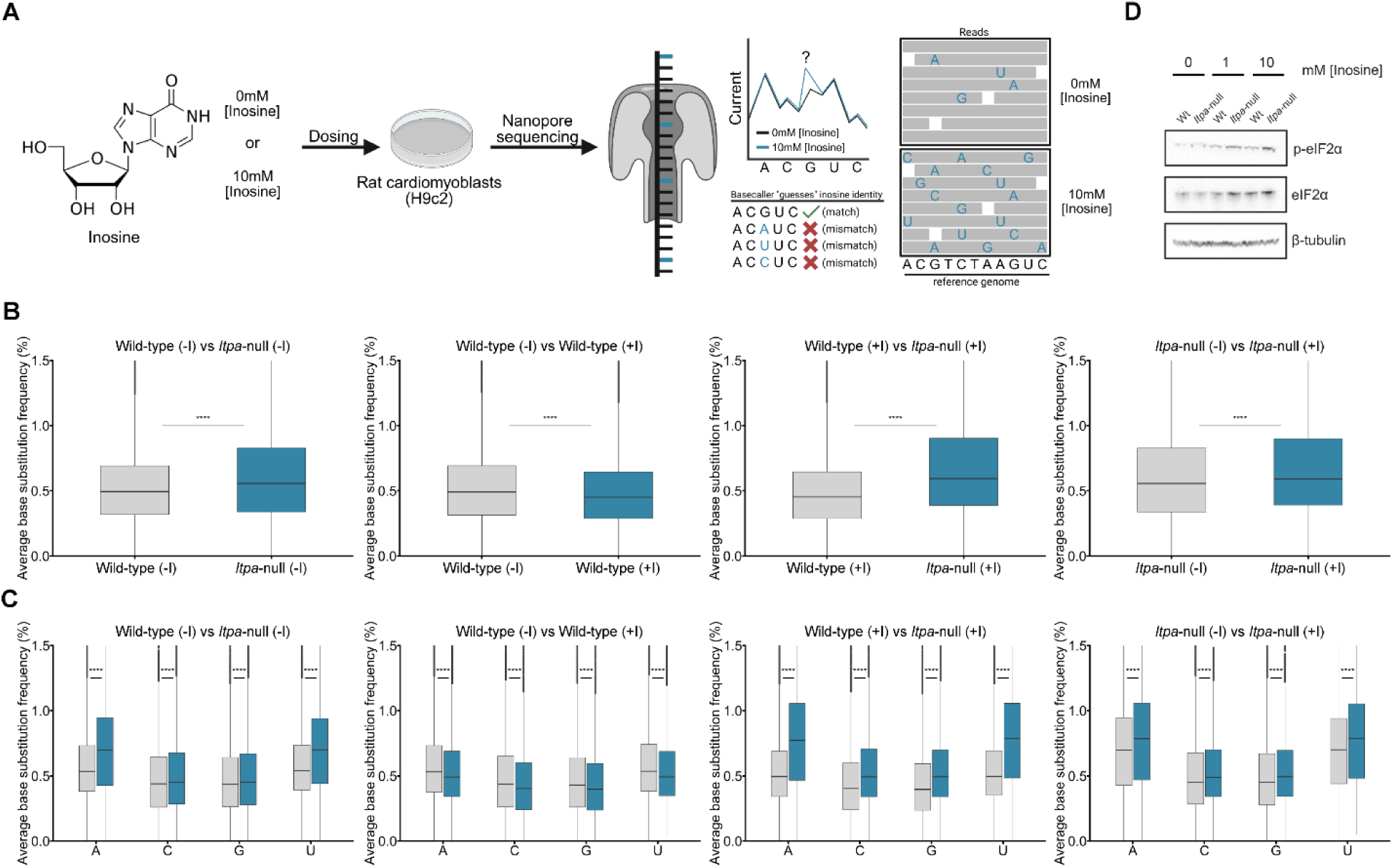
Inosine nucleoside treatment leads to inosine misincorporation into mRNA and activates the ISR in the absence of ITPase enzyme. **(A)** Wild-type and *Itpa*-null H9c2 cells were treated with inosine nucleoside in the cell culture media. RNA was extracted and mRNA was sequenced using Oxford Nanopore Technologies direct RNA sequencing. Inosine misincorporation results in an altered current from the canonical bases as the RNA translocates through the nanopore resulting in increased base miscalling. **(B)** The average base substitution frequency as a percentage of all filtered reads across the mRNA transcriptome for each genotype with or without inosine treatment (-/+ I). **(C)** Overall base substitution frequency for each base of the filtered reads. Student’s t-test, ****P < 0.0001. **(D)** Immunoblot of (phospho) p-eIF2α, eIF2α and β-tubulin as loading control following treatment of wild-type and *Itpa*-null H9c2 cells with 0, 1 and 10 mM inosine nucleoside in the culture media for 24 hours.

Following ONT direct RNA sequencing, base substitution frequencies were calculated from transcript sequence windows spanning a minimum 500 nucleotide coverage and at least 10x read depth present in each genotype and treatment comparison. This yielded between ∼ 7400 – 9800 sequence windows from all 22 chromosomes of the rat genome (Table S1). Comparing overall base substitution frequencies in the untreated (-I) wild-type vs *Itpa*-null samples revealed a significant median increase from 0.5 to 0.58 reflecting elevated inosine misincorporation in the *Itpa*-null mRNA (Figure 5B, Table S2). When inosine nucleoside-treated (+I) and untreated wild-type samples were compared, there was no increase in the overall base substitution rates, consistent with functional ITPase enzyme restricting the misincorporation of inosine into mRNA. In contrast, samples from treated *Itpa*-null cells revealed significant median increases in base-substitution frequency when compared to untreated *Itpa*-null cells (0.62 vs 0.58%); the increase when compared to wild-type treated samples was even greater (0.62 vs 0.46%) (Figure 5B, Table S2). These same trends were evident when evaluating base substitution frequencies at the level of each individual base (Figure 5C, Table S2). Our results are consistent with stochastic misincorporation of inosine in place of any nucleotide within mRNA of *Itpa*-null cells, a situation that is exacerbated with inosine nucleoside treatment.

To confirm activation of the ISR, we performed immunoblotting for phospho-eIF2α in both the wild-type and *Itpa-*null cells treated with increasing inosine nucleoside in the culture media (Figure 5D). While there was low basal phospho-eIF2α signal in ITPase-null cells, likely due to the low inosine levels in the cultured cells, there was elevated phospho-eIF2α signal upon inosine nucleoside treatment. Our data are consistent with the inosine nucleoside treatment resulting in elevated inosine misincorporation into mRNA in the absence of ITPase enzyme, and thereby leading to activation of the ISR.

### *ITPA*-null neurons upregulate stress response genes upon differentiation

Our results implicate inosine misincorporation into mRNA as a potential pathogenic mechanism underlying ITPase deficiency. However, the rate of inosine misincorporation into mRNA in *Itpa*-null H9c2 cells (Figure S7) is lower than that previously observed in heart and brain tissue of *Itpa*-null mice^11,12,15^. Additionally, ITPase-deficient mice appear asymptomatic as embryos but display perinatal lethality with symptoms of dilated cardiomyopathy and audiogenic seizures^11,12,15^, suggesting that developmental timing contributes to pathogenesis. Heart and brain development, involving the generation of a large proportion of postmitotic cells, may be particularly vulnerable to inosine accumulation. To investigate the stress response associated with inosine accumulation in a post-mitotic developmental cell model, we generated *ITPA*-null human iPSCs and then differentiated these into neurons. CRISPR/Cas genome engineering was performed as described in Ran *et al.*, 2013^43^ using vectors for the expression of Cas9, and gRNA targeting exon 4 of *ITPA*. Biallelic frameshift in/dels and unedited alleles to serve as wildtype clones were confirmed through Sanger sequencing (Figure S8). Isogenic wild-type and *ITPA*-null iPSC clonal cell lines were generated in parallel. Western blotting confirms that ITPA protein is absent in the null clones (Figure 6A).

**Figure 6.**
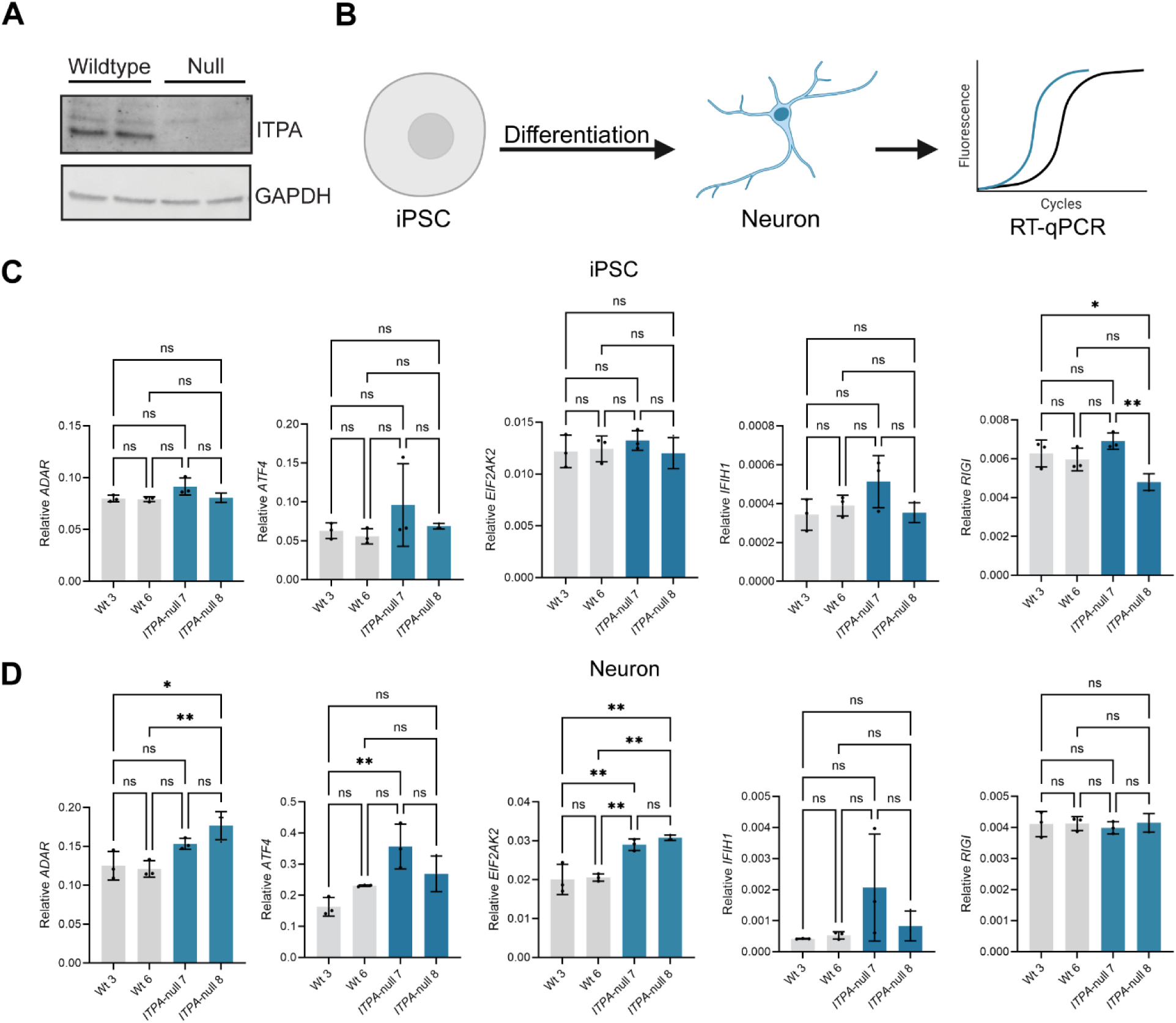
Absence of ITPase enzyme in iPSCs induces low-level stress response gene expression upon differentiation into neurons. **(A)** Immunoblot for ITPA protein in wild-type and *ITPA*-null iPSCs with *GAPDH* as a loading control. **(B)** iPSCs were differentiated into neurons and gene expression was evaluated using RT-qPCR. **(C)** Expression of *ADAR*, *ATF4, EIF2AK2*, *IFIH1* & *RIGI* transcripts relative to *GAPDH* in wild-type and *ITPA*-null iPSCs and **(D)** in differentiated neurons. Mean +/- SD, n = 3 experimental replicates, One-way ANOVA/Tukey’s multiple comparisons, *P < 0.05, **P < 0.01, ns – not significant.

iPSC clones were induced into forebrain neural progenitor cells and matured into neurons over three weeks. Neuronal identity was confirmed by the markers βIII-tubulin and MAP2 (Figure S9) and total RNA was collected from the cultures. Inosine levels were measured using LC-MS. Unexpectedly, we could not detect inosine misincorporation into RNA by these means, perhaps suggesting that inosine levels were lower than the detection limit of our assay. We evaluated expression of our stress response gene panel in both iPSCs and the differentiated neurons (Figure 6B). While there was no significant upregulation of these factors in the *ITPA*-null iPSCs, there was significant upregulation of *ADAR*, *ATF4* and *EIF2AK2* in one or both of the *ITPA*-null clones relative to wild-types upon neuronal differentiation, suggestive of a low-level stress response in the absence of ITPase enzyme (Figure 6C, 6D).

## DISCUSSION

Our study reveals that inosine misincorporation into mRNA is an unappreciated cause of cellular stress. Under normal circumstances, the ITPase enzyme prevents misincorporation of inosine nucleotides into RNA by hydrolyzing ITP, thereby suppressing this form of stress. However, in cells lacking ITPase, we demonstrate that inosine misincorporation into mRNA causes activation of the ISR. Our experiments with transfection of exogenous inosine-containing mRNA into cells underscores this finding, showing even more pronounced stress responses. We speculate that such stress responses contribute to the pathogenesis of ITPase deficiency.

Inosine serves an essential role in mRNA, but is normally selectively introduced through A-to-I editing mediated by the adenosine deaminase acting on RNA (ADAR) family of proteins^25^. Introduction of inosine in a site-specific manner results in protein recoding during translation, while hyper-editing of repetitive sequences including *Alu* elements in non-coding regions of mRNA serves to modulate RNA structure to distinguish “self” from “non-self” RNA and prevent inappropriate innate immune activation^25,44^. Biallelic pathogenic variants in *ADAR* cause the inflammatory disorder Aicardi-Goutières Syndrome (AGS), which is associated with widespread upregulation of interferon stimulated genes^45^. It is interesting that in the context of AGS, inosine in RNA acts as an immune-suppressive signal whereas our findings show that it can also be immune-stimulatory (Figure 1). This suggests that the quantity of inosine in mRNA, and/or its location within mRNA, are important determinants of the effect that it has in cells.

Transfection of inosine-containing mRNA into cells (Figures 2G, S4D, S5), and inosine nucleoside supplementation of ITPase-deficient cells, which results in inosine misincorporation into endogenous mRNA (Figure 5D), both activate the ISR. These findings indicate that a major cellular function of the ITPase enzyme is to restrict inosine misincorporation into mRNA and prevent activation of the ISR. Our results implicate PKR as a potential sensor of inosine misincorporation into mRNA, since inosine-containing mRNA activates phosphorylation of PKR (Figure S4D). Consistent with this finding, PKR inhibition mitigates the stress response to inosine mRNA and rescues cellular viability (Figure 3). Because PKR is stimulated by double-stranded RNA, one potential explanation for inosine misincorporation into mRNA triggering the ISR is that inosine induces double-stranded RNA formation. However, RNA lacking an mRNA cap and polyA tail (Figure S6) or a non-canonical IRES RNA translation substrate (Figure 4E-F), both abrogated the cellular response to inosine misincorporation, indicating that double-stranded RNA formation alone, due to inosine misincorporation, was not sufficient to induce the stress response without the context of a capped and polyadenylated RNA substrate.

Furthermore, the lack of a canonical start codon which severely reduced functional translation, unexpectedly had no effect on the stress response (Figure 4). Therefore, it is also unlikely that inosine misincorporation into mRNA is driving the stress response through the production of aberrant peptides resulting from miscoding during translation or as a result of ribosome collisions/stalling. An alternative mechanism should also account for the stress granule formation we observed in response to inosine-containing mRNA (Figures 2E, 2F). Collectively, these findings raise the intriguing possibility that inosine misincorporation into mRNA activates a unique mRNA stress pathway involving the ribosome, which leads to PKR-mediated activation of the ISR and stress granule formation. Future work to further delineate the precise mechanism of activation will be of considerable interest to expand our understanding of inosine RNA biology and mRNA quality control, particularly given its relevance to diseases, including those caused by ITPase deficiency and perturbed A-to-I editing.

It remains to be seen if inosine misincorporation into mRNA triggers innate immune activation and the ISR *in vivo*. Notably, ITPase-deficient *Arabidopsis thaliana* plants show upregulation of genes associated with biotic stress and systemic acquired resistance, akin to plant innate immunity^13^. Together with our findings, this raises the question of whether ITPase deficiency is associated with the ISR and innate immunity in patients, a question that warrants future investigation. Although we were unable to detect inosine in RNA from *ITPA*-null iPSCs that were differentiated into neurons, there was a low-level pattern of gene expression indicative of cellular stress (Figure 6). It is currently unclear why cultured cells exhibit lower inosine misincorporation rates than those observed in tissues of *Itpa*-null mice and plants^11–13,15^. One possible explanation is that cultured cells rely less on endogenous purine nucleotide synthesis, particularly if canonical nucleotides were present in the culture media, thus leading to diluted inosine accumulation. Another plausible explanation is that ITP accumulation is greater in tissues due to elevated deamination of ATP over time, for example, as a result of higher oxidative stress. Consistent with this latter hypothesis, ITPase-deficient *Arabidopsis thaliana* show elevated inosine misincorporation into RNA as a result of oxidative stress and greater misincorporation in aged versus younger plants^13^. Future work is necessary to determine the relative contributions of these and potentially additional factors to inosine misincorporation rates, and the resulting cellular stress responses.

In summary, our findings suggest that a major cellular role of the ITPase enzyme may be to avert the ISR by restricting inosine misincorporation into mRNA. In the absence of ITPase, targeting ISR activation using PKR inhibitors, poses a potential therapeutic avenue for treating this rare but fatal condition, which warrants future investigation.

## METHODS

### Plasmids

Firefly and *Renilla* luciferase T7 pcDNA3 constructs were generated as described previously^29^. Alternative start codon constructs were generated with site-directed mutagenesis of the Firefly luciferase construct using the Q5 Site-Directed Mutagenesis Kit (NEB). Primers for the site-directed mutagenesis were designed using NEBaseChanger and ordered through IDT (Table S3). All plasmids were sequence verified using Plasmidsaurus service.

### Primers

Primers for RT-qPCR were designed using the IDT PrimerQuest tool and ordered through IDT. Primers for the site-directed mutagenesis were designed using NEBaseChanger and ordered through IDT. All primer sequences are listed in Table S3.

### Cell lines

H9c2 wild-type and *Itpa*-null rat cardiomyoblast cells^15^ were cultured in Dulbecco’s modified Eagle’s medium (DMEM; ThermoFisher) supplemented with 10% fetal bovine serum (FBS) and 1% Gibco Antibiotic-Antimycotic at 37°C and 5% CO_2_. A549 cells were cultured in Dulbecco’s modified Eagle’s medium (DMEM; ThermoFisher) supplemented with 10% fetal bovine serum (FBS), 1% Gibco MEM Non-Essential Amino Acids, 1% Gibco L-Glutamine (200 mM), and 1% Gibco Antibiotic-Antimycotic at 37°C and 5% CO_2_.

The 680.2B-iPSC cell line was provided by the Stem Cell Research Facility at Memorial Sloan Kettering Cancer Center. This line was previously generated from fibroblast cells of a healthy donor purchased from Coriell Institute (GM01680) and has been fully characterized^46^. iPSCs were cultured on Matrigel (Fisher Scientific 08-774-552) in Stemflex Medium (Thermo Fisher A3349401) and maintained at 37 °C with 5% CO2. For regular passaging, cells were detached and passaged with 0.5 mM EDTA (Fisher Scientific MT-46034CI) at room temperature for 7 min.

#### Wild-type and *ITPA*-null iPSC maintenance

All iPSC lines were received to Neuracell Core Facility and screened for mycoplasma contamination. iPSCs were maintained in six-well plates coated with growth factor-reduced Matrigel or Cultrex at 37ᵒC and 5% CO2. The cultures were fed with mTeSR1 medium with an FGF2-DISC (Stem Cultures, DSC500-48). The medium, but not the FGF2-DISC, was replaced every 2-3 days based on culture confluency. iPSC cultures were clump-passaged using ReLeSR (STEMCELL Technologies) about once a week. Cells were not allowed to grow past 80% confluency.

#### Wild-type and *ITPA*-null Neuronal differentiation

Induced pluripotent stem cell (iPSC) colonies were dissociated into single cells using Accutase (Invitrogen) for 8 minutes at 37°C. Cells were washed twice with DMEM/F12 (Invitrogen) and replated onto plates coated with Laminin 521 (Corning, Cat# 354222). Initial seeding was performed in mTeSR medium supplemented with a Rho-associated protein kinase (ROCK) inhibitor. Forebrain neural induction was initiated 24 hours post-seeding (Day 0) using a dual SMAD inhibition with minor modifications^47^. Neural progenitor cells (NPCs) were harvested and cryopreserved on day 20–21. For subsequent experiments, thawed NPCs were plated at high density on Laminin 521 and maintained in NM medium with daily changes for 5–7 days until confluent and replated for terminal neuronal differentiation. NPCs were dissociated with Accutase and seeded at a density of 100,000 cells/cm². Cell viability and density were determined using a hemocytometer and Trypan Blue exclusion (Sigma-Aldrich). Cells were plated onto tissue culture-treated plates coated with Laminin 521. Differentiation was conducted in BrainPhys medium supplemented with 1% B27 with vitamin A (Life Technologies), 10 ng/mL BDNF, and 10 ng/mL GDNF. Neuronal cultures were maintained for three weeks with medium changes performed three times per week.

### Inosine nucleoside treatment

H9c2 cells were seeded in 6-well plates at 500000 cells/well and allowed to adhere over 24 hours. Inosine nucleoside was dissolved in DMSO and added to cells in the media to final concentrations of 0, 1 and 10 mM. At the time of harvest, media was removed, and cells were washed in 1 mL of 1x PBS.

### LC-MS analysis of inosine and adenosine in RNA

#### RNA sample preparation

500 ng of RNA extracted from cells (Aurum^TM^ Total RNA Mini Kit, Bio-Rad) was digested with Nucleoside Digestion Mix (New England BioLabs) according to the manufacturer’s instructions. Next, 60ul of LC-MS grade acetonitrile (A955, Fisher Scientific) was added to 20ul of digested RNA and incubated on ice for 30 minutes with vortexing. Following incubation, samples were centrifuged at 17,000x g, 4 degrees Celsius, for 10 minutes and supernatant was transferred to glass vials for LC-MS analysis.

#### LC/MS analysis

Samples were analyzed by high resolution mass spectrometry with an Orbitrap Exploris 240 (Thermo) coupled to a Vanquish Flex liquid chromatography system (Thermo). 5 µL of samples were injected on a Waters XBridge XP BEH Amide column (150 mm length × 2.1 mm id, 2.5 µm particle size) maintained at 25C, with a Waters XBridge XP VanGuard BEH Amide (5 mm × 2.1 mm id, 2.5 µm particle size) guard column. For positive mode acquisition, mobile phase A was 100% LC-MS grade H2O with 10 mM ammonium formate and 0.125% formic acid. Mobile phase B was 90% acetonitrile with 10 mM ammonium formate and 0.125% formic acid. The gradient was 0 minutes, 100% B; 2 minutes, 100% B; 3 minutes, 90% B; 5 minutes, 90% B; 6 minutes, 85% B; 7 minutes, 85% B; 8 minutes, 75% B; 9 minutes, 75% B; 10 minutes, 55% B; 12 minutes, 55% B; 13 minutes, 35%, 20 minutes, 35% B; 20.1 minutes, 35% B; 20.6 minutes, 100% B; 22.2 minutes, 100% B all at a flow rate of 150 μl min−1, followed by 22.7 minutes, 100% B; 27.9 minutes, 100% B at a flow rate of 300 μl min−1, and finally 28 minutes, 100% B at flow rate of 150 μl min−1, for a total length of 28 minutes. The H-ESI source was operated in positive mode at spray voltage 3500 with the following parameters: sheath gas 35 au, aux gas 7 au, sweep gas 0 au, ion transfer tube temperature 320 C, vaporizer temperature 275 C, mass range 260 to 800 m/z, MS1 mass resolution of 120,000 FWHM, RF lens at 70%, standard automatic gain control (AGC), MS2 fragmentation with normalized collision energies of 20%, 30%, 50%, 75%, 100% and resolution at 15,000 FWHM. LC-MS data were analyzed by Maven software for peak area determination and annotation via matching to external standards. Calibration curves were plotted using linear regression.

### T7 *in vitro* transcription

T7 *in vitro* transcription reactions were carried out as described previously^29^. Briefly, Firefly and Renilla luciferase constructs were linearized using NEB EcoRV-HF or Afe1 in rCutSmart buffer and gel-extracted (NEB Monarch DNA Gel Extraction Kit #T1120S) according to the manufacturer’s instructions. In vitro transcription (IVT) reactions were carried out using the HiScribe T7 ARCA mRNA Kit (with tailing) (NEB) in all capped, polyadenylated experiments. The HiScribe T7 Quick High Yield RNA Synthesis Kit (NEB) was used for IVT of uncapped, non-polyadenylated RNA as well as the HCV-IRES-FLuc RNA. IVT reactions were carried out in a 20 μl volume with 100 ng of DNA template in the presence of 10 mM of each canonical nucleotide. ITP (Sigma-Aldrich 10879) was added to a final concentration of 0, 0.1 or 1 mM. Reactions were incubated at 37°C for 1 h, then treated with DNase I for an additional 15 min. RNAs were column-purified (NEB Monarch RNA Cleanup Kit #T2040L), eluted in nuclease-free H_2_O and stored at –80°C.

### Luciferase assay following mRNA transfections

H9c2 and A549 cells were seeded in 96-well plates at 10000 cells/wells for 24 hours. RNA (12.5 ng) was diluted in 5 µL of Gibco Opti-MEM. Invitrogen MessengerMAX (0.5 µL) and 5 µL of Opti-MEM were incubated for 10 minutes at room temperature. The 5.5 µL mix was added to the RNA in Opti-MEM and incubated for 5 minutes at room temperature. The final 10.5 µL transfection mix was added to each well. Cells were harvested between 24-72hr as stated with media change at 24 hours post-transfection. Cells were lysed in Promega Cell Culture 5X Lysis reagent diluted to 1X in MBG H_2_O for 5 minutes using pipette mixing and a lab rocker. Cell lysate (20 µL) was transferred to a 96-well assay plate. A Promega Glomax 96 microplate luminometer was set for 40 µL injection of Promega Luciferase Assay Regent (firefly constructs) or the Promega Renilla Luciferase Assay System (renilla construct) for 0.2 second delay and 5 second integration time. Raw luciferase measurements were normalized to the control RNA in all experiments. Computation of averages, standard deviations and statistics were made in RStudio v2024.09.0+375 and plotted using ggplot2 v3.5.1. For PKR inhibitor experiments, RNA input was increased to 50 ng and at 6 hours post-transfection, media was replaced with fresh media containing 0 (DMSO only) or 1 µM Imidazolo-oxindole PKR inhibitor C16 (Sigma-Aldrich # 527450) and treatment was performed for 24 hours. For A549 HCV-IRES-FLuc RNA experiments, input was increased to 250 ng and transfection time adjusted to 18 hours.

### RT-qPCR following mRNA transfections

Cells were seeded in a 12-well plate at 100000 cells/wells for 24 hours. RNA (125 ng) was diluted in 50 µL of Gibco Opti-MEM. Invitrogen MessengerMAX (2 µL) and 50 µL of Opti-MEM were incubated for 10 minutes at room temperature. The 52 µL mix was added to the RNA in Opti-MEM and incubated for 5 minutes at room temperature. The final 102 µL transfection mix was added to each well. At the time of harvest (24-72hr) as stated, media was removed, cells were washed in 1 mL of 1x PBS. For transfections exceeding 30 hours, media was changed every 24 hours. For PKR inhibitor experiments, at 6 hours post-transfection, media was replaced with fresh media and Imidazolo-oxindole PKR inhibitor C16 inhibitor (Sigma-Aldrich 527450) at a final concentration of 0 (DMSO only) or 1 µM; transfection was extended to 30 hours. For A549 cells transfected with FLuc mRNA, 50ng of input was used; for the HCV-IRES-FLuc, 1250ng input RNA was used and time adjusted to 18 hours.

RNA was extracted from each well using the Bio-Rad Aurum total RNA Mini kit according to the manufacturer’s instructions in a 26 µL elution volume. RNA concentration was measured using a ThermoFisher NanoDrop One UV-Vis spectrophotometer. RNA (∼250ng) was reverse transcribed with 10 µM random hexamer primers followed by reverse transcription with ThermoFisher SuperScript IV reverse transcriptase in a 20 µL final volume. Following RT, RT reaction was diluted 1:2. A 384 or 96-well plate was loaded with 1 µL of cDNA, 1 µL of a forward and reverse primer mix (10 µM each), 3 µL of MBG H_2_O, and 5 µL of ThermoFisher PowerUp SYBR Green Master Mix. Plates were sealed and briefly centrifuged in a microplate spinner.

Plates were run in a Bio-Rad CFX384 or CFX96 Touch RealTime PCR Detection System. ΔΔCq values and averages were calculated in Microsoft Excel, standard deviations and statistics were computed in RStudio v2024.09.0+375 and plotted using ggplot2 v3.5.1 or using Graphpad Prism 10.

### Cell viability assay

Cells were seeded in a 96-well plate at 10000 cells/wells for 24 hours. RNA (50 ng) was diluted in 5 µL of Gibco Opti-MEM. Invitrogen MessengerMAX (0.5 µL) and 5 µL of Opti-MEM were incubated for 10 minutes at room temperature. The 5.5 µL mix was added to the RNA in Opti-MEM and incubated for 5 minutes at room temperature. The final 10.5 µL transfection mix was added to each well. For PKR inhibitor treatments, mRNA transfection input was increased to 100 ng and at 6 hours post-transfection, cell culture media was replaced with fresh media containing 0 (DMSO only) or 1 µM Imidazolo-oxindole PKR inhibitor C16 (Sigma-Aldrich). The staurosporine (positive control) was added at a final concentration of 250 nM. Before harvest (21 hours), media was removed, cells were washed in 200 µL of 1x PBS, 50 µL of media was added along with 50 µL of MTT (5 mg/mL). At the time of harvest (24 hours), media was removed and 100 µL of DMSO was added. The 96-well plate was wrapped in foil and placed on an orbital shaker at low-medium speed for 5 minutes. Each well of the 96-well plate was measured for absorbance at 570 nm using a BioTek Synergy H1 Multimode Reader. All treatments were normalized to the 0mM (no inosine) condition. Computation of averages, standard deviations and statistics were made in RStudio and plotted using ggplot2.

### Polysome profiling

Cells were seeded in a 10 cm plate at 1.5 M cells/well for 24 hours. Media was removed and replaced with 8 mL of fresh media. RNA (1.8 µg) was diluted in 1 mL of Gibco Opti-MEM. Invitrogen MessengerMAX (38.5 µL) and 961.5 µL of Opti-MEM were incubated for 10 minutes at room temperature. The 1 mL mix was added to the RNA in Opti-MEM and incubated for 5 minutes at room temperature. The final 2 mL transfection mix was added to each plate now at a total volume of 10 mL. Before harvest (21 hours) media was replaced with 5 mL of fresh media. At the time of harvest (24hr), 50 µL of 10 mg/mL cycloheximide was added and incubated for 3 minutes at 37°C and 5% CO_2_. Media was removed and 5 mL of 4°C 1X PBS was added to sides of plate for gentle washing twice followed by removal by vacuum aspiration. Lysis solution (300 mM NaCl, 15 mM MgCl_2_, 0.1 mg/mL cycloheximide, 0.1 mg/mL heparin, 1% triton X-100) was added to cells (400 µL) and scraped using a cell scraper. Lysate was added to a microfuge tube and incubated for 10 minutes at 4°C. Microfuge tubes were centrifuged for 5 minutes in a 4°C centrifuge at 8400 RPM. A 10-50% sucrose gradient was made using 10% sucrose solution (10% sucrose, 0.1 mg/mL cycloheximide, 300 mM NaCl, 15 mM MgCl_2_) and 50% sucrose solution (50% sucrose, 0.1 mg/mL cycloheximide, 300 mM NaCl, 15 mM MgCl_2_) with a Biocomp Gradient Master 108 in a 14 X 89 mm tube. Supernatant of lysate was loaded onto each gradient and centrifuged at 35000 g, for 2 hours and 35 minutes at 4°C with vacuum in a Beckman Coulter Optima L-90K ultracentrifuge. The Brandel Syn-202 Density Gradient Fractionation system was used to measure absorbance values of the centrifuged gradients with PeakChart software v1.0.0.0. The background absorbance was determined by finding the lowest value in each trace. The background absorbance was subtracted from all raw absorbance values. Area under the curve normalization was performed by calculating the sum of the absorbance values for the whole trace and then dividing each absorbance value by the sum. Polysome signal reductions were calculated using the formula (treatment – control) / control X 100) on the polysome regions of each profile. The 1mM [ITP] FLuc profile values were dilated on the x-axis following the reduction calculations to align the monosome peaks. The normalized absorbance values were used to generate the polysome profiles in RStudio with ggplot2.

### Stress granule detection

Cells were plated at 20000 cells/well in an 8-well chamber slide for 24 hours. RNA (25, 50, or 100 ng) was diluted in 10 µL of Gibco Opti-MEM. Invitrogen MessengerMAX (1 µL) and 9 µL of Opti-MEM were incubated for 10 minutes at room temperature. The 10 µL mix was added to the RNA in Opti-MEM and incubated for 5 minutes at room temperature. The final 20 µL transfection mix was added to each well. At the time of harvest (24 hr), media was removed, cells were washed in 1 mL of 1x PBS. Cells were fixed with 4% paraformaldehyde in PBS for 20 minutes. Cells were washed twice with PBS for 5 minutes. Cells were permeabilized with 0.5% Trion X-100/1% fish gelatine in PBS for 10 minutes. G3BP1 rabbit primary antibody (Thermo Fisher A302-034A; 100 µL) was added to wells and incubated overnight (∼16 hours) at 4°C. Cells were washed twice with 1% fish gelatine in PBS for 10 minutes. Secondary antibody was diluted in 1% fish gelatine in PBS (100 µL final) and added to wells for 1 hour covered at room temperature. Cells were washed twice with 1% fish gelatine in PBS for 5 minutes. Invitrogen Hoechst 33342 (10000 fold dilution) in 1% fish gelatine in PBS was added to cells for 5 minutes. Cells were washed with 1% fish gelatine in PBS for 5 minutes. Fluoromount (5 µL) was added to each well, covered with a coverslip, and secured using nail polish. Stress granules and nuclei were visualized using a Zeiss LSM 980 confocal microscope. Images were acquired with the Plan-Apo 63X/Oil DIC 1.4 NA objective in channel mode to obtain a 2-channel image. Multiple fields of view (n(FOV)=7 per group) were manually selected to provide representative coverage of each sample for quantitative analysis. Analysis was done in ImageJ v1.54f. All the figures and statistics were generated in R.

### Immunoblotting

Cells were seeded at 625000 cells/T25 flask for 24 hours for transfection experiments and at 1000000 cells/10cm dish for inosine nucleoside treatment experiments. In transfection experiments, RNA (625 ng) was diluted in 250 µL of Gibco Opti-MEM. Invitrogen MessengerMAX (10 µL) and 250 µL of Opti-MEM were incubated for 10 minutes at room temperature. The 260 µL mix was added to the RNA in Opti-MEM and incubated for 5 minutes at room temperature. The final 510 µL transfection mix was added to each well. Cells were washed twice with 4°C 1x PBS. In inosine treatment experiments, inosine in DMSO, or DMSO alone, was diluted to the appropriate concentration in media and then these dilutions were used to replace the media on each plate. In both sets of experiments, cells were incubated for 24 hours, and then trypsinized, pelleted and washed twice with PBS. Pellets were lysed with RIPA buffer (GBiosciences, 786-489) containing proteinase and phosphatase inhibitors (Roche, 04693159001 and 4906845001), on ice, and then lysates were clarified by centrifugation. Following protein quantification, clarified lysates were combined with NuPAGE LDS sample buffer (Thermo, NP0007) containing β-mercaptoethanol as a reducing agent. Samples were separated by electrophoresis on 4–12% Bis–Tris gels in 1X MOPS buffer (Biorad, 3450124, 1710788). Protein was transferred to PVDF membranes using a Trans-Blot Turbo device and associated reagents (Biorad, 1704275). Membranes were blocked in 5% milk in 1X TBST for at least 1 hour prior to probing with antibodies. Membranes were probed with anti-eIF2α (1:1000), anti-p-eIF2α (1:1000) (Cell Signaling Technology, 5324 and 3398), anti-PKR (1:1000), anti-p-PKR (1:1000) (AbCam, ab184257 and ab32036, respectively), anti- β-tubulin (1:5000) (Cell Signaling Technology, 2128) or anti-actin (1:1000) (Sigma A2066) overnight at 4°C. Probed membranes were washed three times in 1X TBST then further probed with HRP-conjugated donkey anti-rabbit secondary antibody (1:10000) (ThermoFisher, 31458)for at least 1 hour. Membranes were washed three times in TBST, then developed using ECL Plus Western Blotting Substrate (Pierce, 32132) according to manufacturer’s instructions. Bands were visualized using a Bio-Rad ChemiDoc MP Imaging System.

#### Immunoblotting for wild-type and *ITPA*-null iPSCs

One well of a six-well plate was harvested per iPSC clone, pelleted, and stored at -80. iPSC pellets were thawed on ice and lysed in radioimmunoprecipitation assay (RIPA) buffer (Thermo Scientific) with 1X cOmplete Protease Inhibitors (Roche) for 15 min on ice, centrifuged at 21,000x g for 15 min at 4°C and the supernatant was collected as the protein lysate. Protein lysate concentration was quantified using the Pierce BCA Protein Assay Kit (Thermo Scientific) and 15µg of soluble protein lysate, with NuPAGE™ Sample Reducing Agent and NuPAGE™ LDS Sample Buffer (both Invitrogen) was separated on a 4-12% Bis-Tris gel (Bio-Rad) with XT MOPS running buffer (Bio-Rad). Protein was then transferred to a 0.2µm nitrocellulose membrane (Amersham™ Protran™, cyvita) in 1x NuPAGE™ Transfer Buffer (Invitrogen) with 20% MeOH at 0.3A for 2 hours at 4°C. The membrane was rinsed in deionized water and blocked in 5% dry milk in 1x PBS containing 0.05% Tween-20 (Sigma) for 2 hours at room temperature. The membrane was probed with rabbity polyclonal α-ITPA antibody (1:500, Origene TA350543) and mouse α-GAPDH (1:1000, Abcam ab8245) in blocking solution overnight at 4°C, washed 3 times for 5 minutes each in 1x PBST and incubated for 1 hour at room temperature with goat α-rabbit HRP (1:10000, Invitrogen A16110) and donkey α-mouse IR Dye 680 (LiCor, cat# 926-68072) in blocking solution. Following 3 x 5 minute washes in 1x PBST, ProSignal Pico Spray (Prometheus, Genesee Scientific) was used following the manufacturer’s instructions and membranes were imaged using a ChemiDoc MP Imaging System (Bio-Rad) imaging the IR Dye 680 channel before the chemiluminescent channel.

### Illumina RNA-seq library preparation

The NEBNext Ultra II Directional RNA Library Prep Kit (Illumina) with NEBNext rRNA Depletion Kit were used to prepare RNAseq libraries, with a total of 250 ng of input RNA. The protocol was carried out following the manufacturer’s instructions, with the following exceptions: 40X adaptor dilutions were used; all bead incubations were performed at room temperature; 10 µM concentrations of the index primers were used; and 10 cycles of library amplification were performed. The resulting libraries were quantified using the NEBnext Library Quant kit (Illumina), inspected for quality and read length distribution via capillary electrophoresis on a Fragment Analyzer using the NGS Analysis DNF-474 kit (Advanced Analytical) and sequenced on an Illumina NextSeq 2000 sequencer.

### Illumina RNA-seq data analysis

FASTQ files were aligned to the GRCr8 *Rattus norvegicus* genome and the *Rattus norvegicus* GRCr8 114 GTF annotation file using STAR alignment tool v2.7.1b (quantmode GeneCounts)^48^ to produce gene-level counts outputs. RStudio was used for processing gene counts data. DESeq2 v1.44.0^30^ was used to perform differential gene expression analysis. Significance was determined using DESeq2 computed log_2_ fold changes and p-adjusted values (log_2_ fold ≥ 1.5, p-adj < 0.05). The R packages ggplot2 v3.5.1 and ggpubr v0.60 were used for data visualizations. Gene ontology analysis was performed using PANTHER v19.0^49^ using the overrepresentation test selected for GO biological process.

### Oxford Nanopore direct RNA sequencing library preparation

The ONT Direct RNA Sequencing (SQK-RNA004) protocol version DRS_9195_v4_revI_30Jul2025 was used to generate libraries. Approximately 1 µg of total RNA sample along with 1 µL of diluted RNA CS (calibrant strand) was ligated to RT Adapter (RTA) with T4 DNA Ligase 2M U/ml (NEB M0202M) and reverse-transcribed (Induro® Reverse Transcriptase and 5x Induro® RT Reaction Buffer), then purified using 1.8× RNAClean XP beads (Beckman 131 Coulter® A63987), with 70% ethanol washes, on a magnetic stand. RNA Ligation Adapter (RLA) was ligated and products were purified using 1× RNAClean XP beads, with 2× WSB wash steps. The final library was eluted in RNA elution buffer (REB) and mixed with Sequencing Buffer (SB) and Library Solution (LIS). Freshly prepared libraries were immediately loaded onto a FLO-PRO004RA flow cell and sequenced using a PromethION sequencer and MinKNOW software v25.05.14.

### Oxford Nanopore direct RNA sequencing analysis

Dorado v1.1.1 was used to basecall the raw pod5 files producing bam file output using the dorado [basecaller] with the parameters sup,inosine_m6A_20meA. Dorado [aligner] with the options "-x splice -k 14" was used to align reads to the rat reference genome GRCr8 producing BAM alignment files. SAMtools v1.13 was used to sort and index BAM files. SAMtools [depth] was used to find genomic loci with ≥10 depth. The depth files were reordered lexicographically using the Bash command sort. A composite key was created combining the chromosome and position columns into the format (chr:pos). The composite key files were joined using the Bash command join to find regions of shared depth between the compared libraries. The composite keys in the joined output file were separated back into separate chromosome and position columns. The separated joined file was converted to BED format and sorted lexicographically. BEDtools [merge] v2.31.1 was used to merge adjacent regions into intervals as a new BED file. The Bash command awk was used to filter for intervals of ≥500 bp in length and create a file containing the genomic windows as coordinates (chr:start:end). The genomic windows-containing file was parsed by a loop that iterated through each line inputting the coordinates into Pysamstats v1.1.2 [variation-strand] to produce statistics output files containing information of matches and mismatches for every position within each genomic window. RStudio was used to calculate the base substitution frequency [mismatches/(matches+mismatches) * 100] for each position within each window. An average base substitution frequency was calculated for each of the 4 nucleotides producing four averages, one for each nucleotide in each window. The base substitution frequencies were plotted using ggplot v3.5.1 in RStudio. For the combined base substitution frequency figures, the averages from each nucleotide were plotted together as part of a single group. A Wilcoxon signed-rank test was used to determine if there was a significant difference in the medians of each library comparison. The R packages GenomicRanges v1.56.2 and rtracklayer v1.64.0 were used to cross reference the window positions with gene annotations in the *Rattus norvegicus* GRCr8 114 GTF annotation file to determine which genes corresponded to each window analyzed.

## ACKNOWLEDGMENTS

We thank J. Cleary from the RNA Institute, UAlbany for assistance with figure preparation. All figure schematics were generated using BioRender. This work was supported in part by the University of Rochester Medical Center (URMC) Metabolomics Resource. We are grateful to the NeuraCell Core Facility team at Neural Stem Cell Institute for their contributions to the iPSC culture and neuronal differentiations. We appreciate the support of the New York State Department of Economic Development’s Center of Excellence in RNA Research and Therapeutics (CERRT).

## AUTHOR CONTRIBUTIONS

K.R., G.F. and M.T.H. conceptualized the study; S.T., A.M.V., C.T.P., J.A.B., K.R., G.F. and M.T.H. supervised the research and interpreted data; J.H.S., N.N.A., E.E.A., M.A.B., H.K.S., B.R.S., T.Z., S.L., T.B. and M.T.H. performed experiments and interpreted data; J.H.S and K.R. wrote the manuscript with input from all authors.

## DATA AVAILABILITY

The Illumina RNA sequencing and Oxford Nanopore Direct RNA sequencing data (BioProject ID: PRJNA1347087) have been deposited in the Sequence Read Archive (SRA) database, https://www.ncbi.nlm.nih.gov/sra.

