## Supplemental Material for "Inosine misincorporation into mRNA triggers the integrated stress response and activates an innate immune gene expression signature"

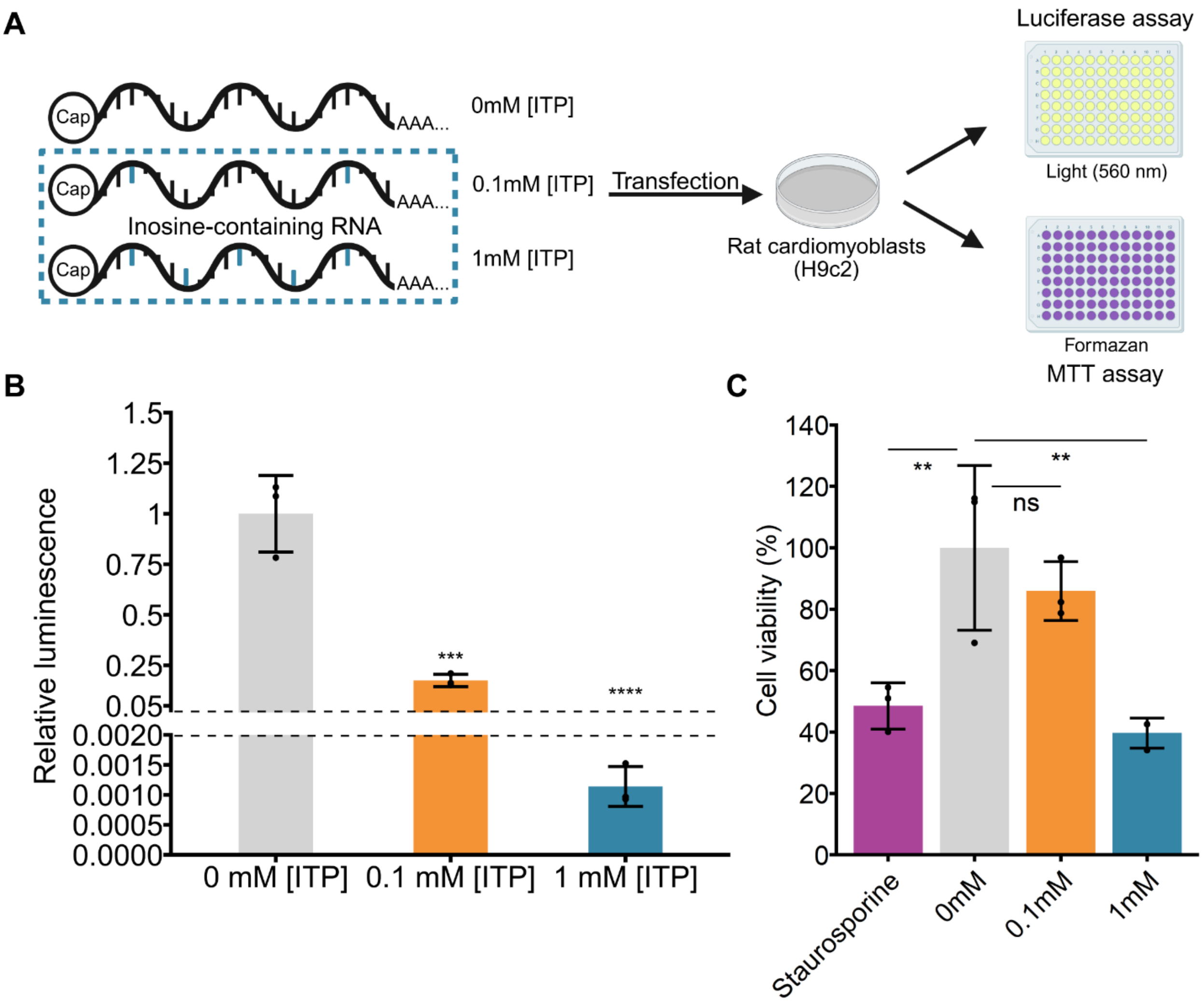


**Figure S1. (A)** Capped firefly luciferase mRNA was *in vitro* transcribed with 0, 0.1 or 1 mM inosine triphosphate [ITP] in the reaction, polyadenylated and then transfected into H9c2 rat cardiomyoblast cells for 24 hours and then luciferase and MTT assays were performed to measure luminescence and cell viability, respectively. **(B)** Luminescence relative to 0mM [ITP] control mRNA. Mean +/- SD, n = 3 experimental replicates, one-way ANOVA/Dunnett's test, ***P < 0.001, ****P < 0.0001. **(C)** Cell viability is expressed as a percentage relative to 0 mM [ITP] control mRNA. Staurosporine treatment is included as a positive control. Mean +/- SD, n = 3 experimental replicates, One-way ANOVA and Tukey's HSD, **P < 0.01, ***P < 0.001, ****P < 0.0001, ns – not significant.

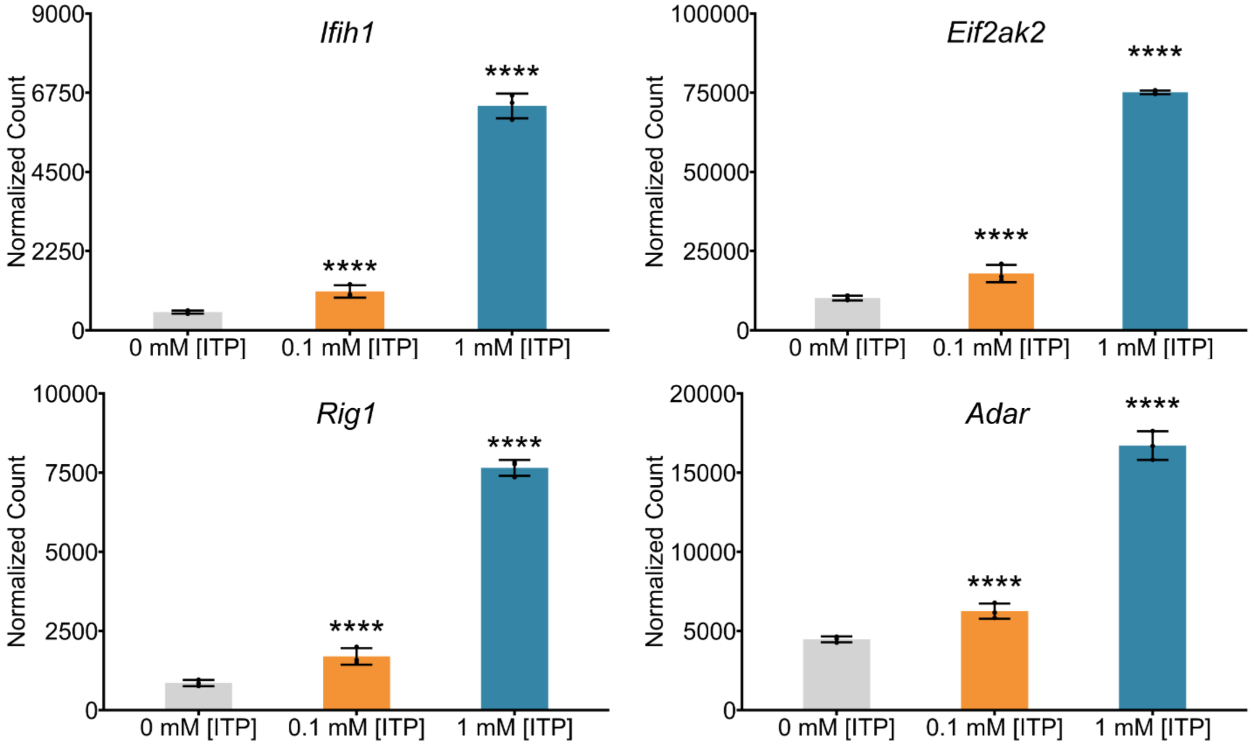


**Figure S2.** DESeq2 normalized read counts of innate immunity genes from Illumina RNAseq of H9c2 rat cardiomyoblast cells transfected with 0, 0.1 and 1 mM [ITP] firefly luciferase RNA for 24 hours, ****P < 0.0001.

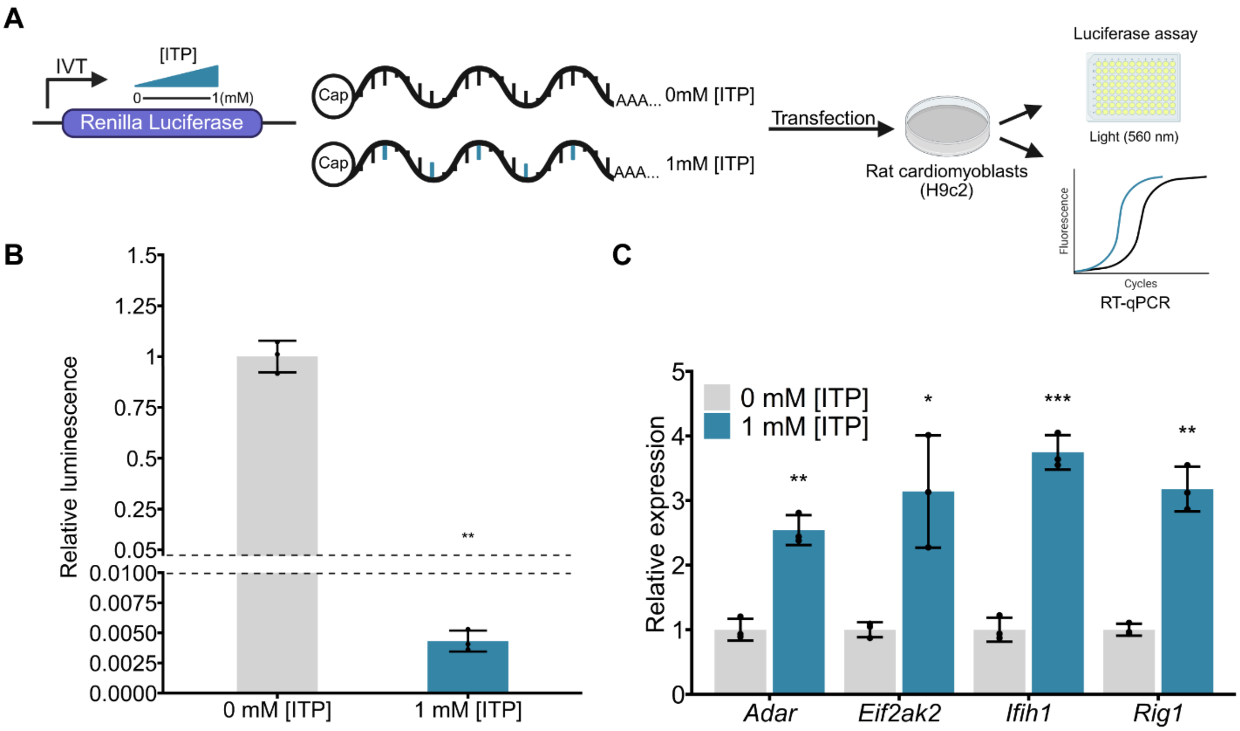


**Figure S3. (A)** Capped *Renilla* luciferase (RLuc) mRNA was *in vitro* transcribed with 0 or 1 mM inosine triphosphate [ITP] in the reaction, polyadenylated and then transfected into H9c2 rat cardiomyoblast cells for 30 hours and then luciferase assay and RT-qPCR were performed. **(B)** Luminescence relative to 0mM [ITP] control RLuc mRNA. Mean +/- SD, n = 3 experimental replicates, Student's t-test, **P < 0.01. **(C)** Expression of the indicated innate immunity genes relative to *Gapdh* and normalized to 0 mM [ITP] control RLuc mRNA for each factor. Mean +/- SD, n = 3 experimental replicates, Student's t-test with Holm-Bonferroni correction, *P < 0.05, **P < 0.01, ***P < 0.001.

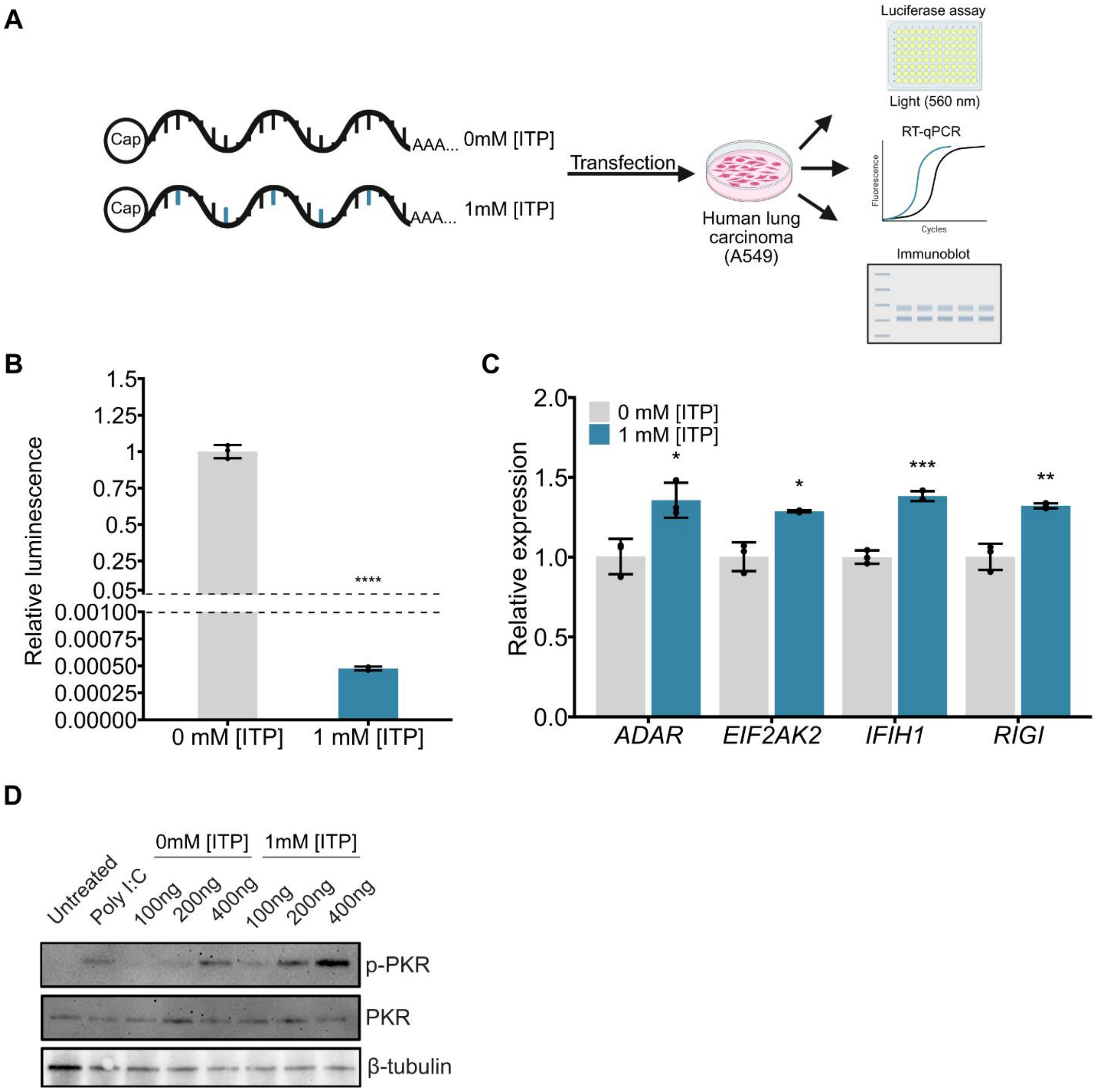


**Figure S4. (A)** Capped firefly luciferase (FLuc) mRNA was *in vitro* transcribed with 0, or 1 mM inosine triphosphate [ITP] in the reaction, polyadenylated and then transfected into A549 human lung carcinoma cells for 18 hours, followed by luciferase assay for luminescence, RT-qPCR and immunoblotting. **(B)** Luminescence relative to 0 mM [ITP] control FLuc mRNA. Mean +/- SD, n = 3 experimental replicates, Student's t-test, ****P < 0.0001. **(C)** Expression of the indicated innate immunity genes relative to *Gapdh* and normalized to 0 mM [ITP] control FLuc mRNA for each factor. Mean +/- SD, n = 3 experimental replicates, Student's t-test with Holm-Bonferroni correction, *P < 0.05, **P < 0.01, ***P < 0.001. **(D)** Immunoblot of (phospho) p-PKR, PKR and β-tubulin as a loading control following transfection of the indicated amount of mRNA. The synthetic double stranded RNA mimetic Poly I:C (100ng) was included as a positive control for p-PKR.

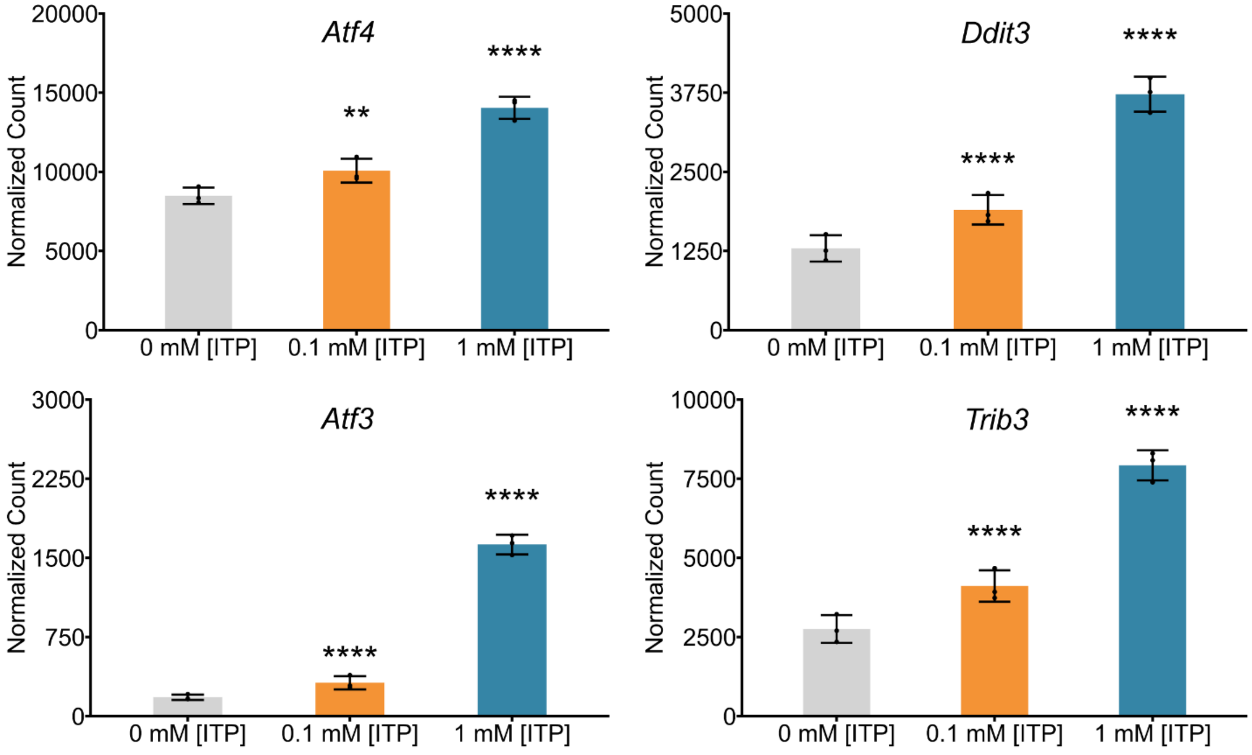


**Figure S5.** DESeq2 normalized read counts of integrated stress response genes from Illumina RNAseq of H9c2 rat cardiomyoblast cells transfected with 0, 0.1, and 1 mM [ITP] firefly luciferase RNA for 24 hours, **P < 0.01, ****P < 0.0001.

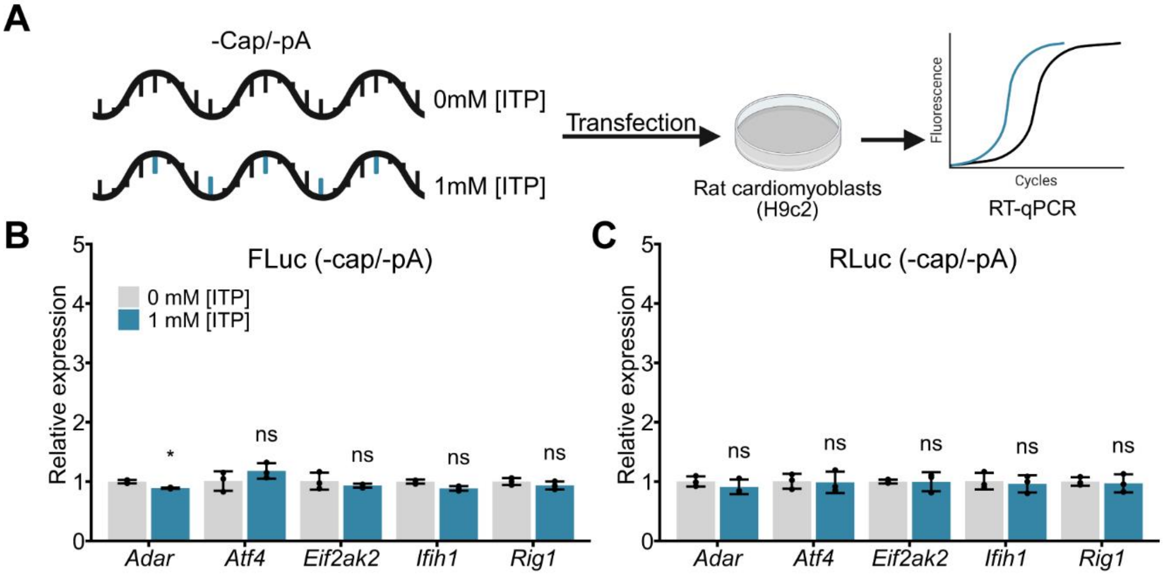


**Figure S6. (A)** Firefly luciferase (FLuc) and *Renilla* luciferase (RLuc) RNA lacking a cap or polyA tail was *in vitro* transcribed with 0 or 1 mM inosine triphosphate [ITP] in the reaction and then transfected into H9c2 rat cardiomyoblast cells for 30 hours, followed by RT-qPCR. Expression of the indicated genes relative to *Gapdh* and normalized to 0 mM [ITP] control RNA for transfections of **(B)** FLuc RNA and **(C)** RLuc RNA. Mean +/- SD, n = 3 experimental replicates, Student's t-test with Holm-Bonferroni correction, *P < 0.05, ns – not significant.

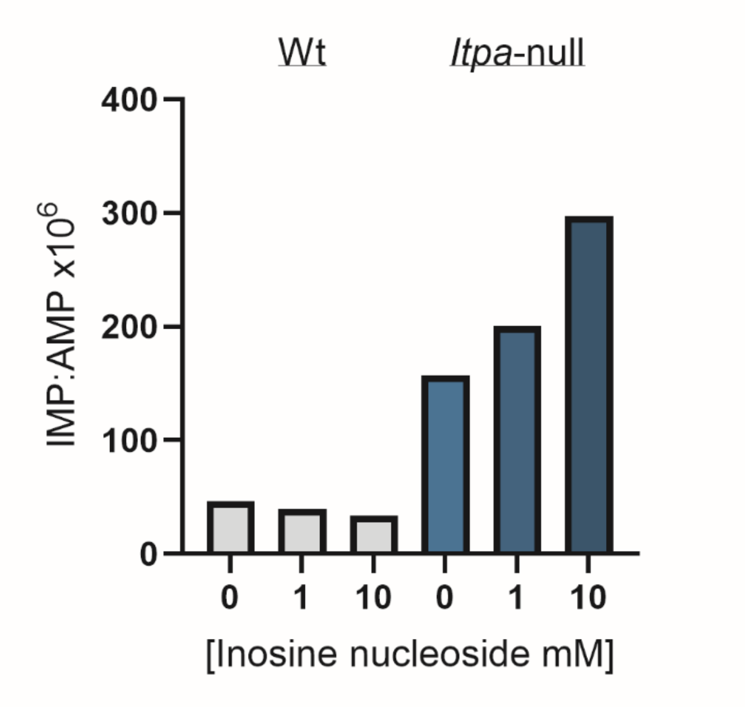


**Figure S7.** Inosine content in total RNA preparations from wild-type (Wt) and *Itpa*-null cells treated with the indicated concentrations of inosine nucleoside in the cell culture media. Total cellular RNA was extracted and digested into nucleosides, and inosine and adenosine levels were quantified using mass spectrometry.

| Gene | sgRNA target | PCR-F1 | PCR-R1 | PCR-R2 (used for seq) |
|---|---|---|---|---|
| ITPA KO in 680.2B iPSCs | ACAAGTGTCCTCAACC AGCA | agtgtgaatcccagctctcc | GGACAACGTGAAAG GCTGTT | CGTGAAAGGCTGTT GATGCT |

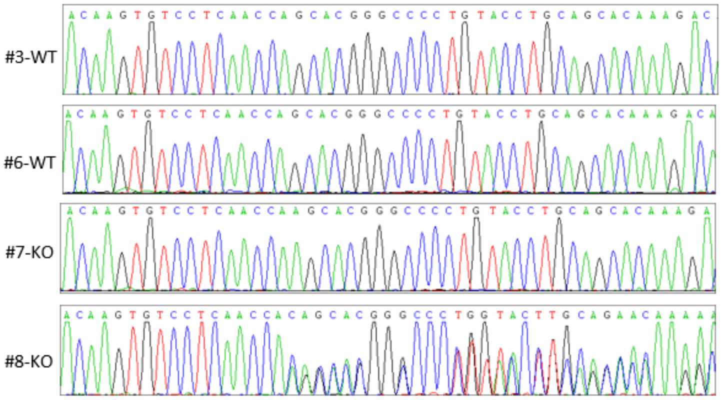


(3; 6) WT : ACAAGTGTCCTCAACCAGCACGGGCCCCTGTACCTGCAGCACAAAGACA

(7) KO: ACAAGTGTCCTCAACCAAGCACGGGCCCCTGTACCTGCAGCACAAAGACA (Insert 1/2; homozygous)

(8) KO-allele 1: ACAAGTGTCCTCAACCAAGCACGGGCCCCTGTACCTGCAGCACAAAGAC (Insert 1)
-allele 2: ACAAGTGTCCTCAACCACAGCACGGGCCCCTGTACCTGCAGCACAAAGA (Insert 2)

**Figure S8.** Sanger sequencing confirms wild-type and *ITPA*-null genotypes of iPSC clonal lines.

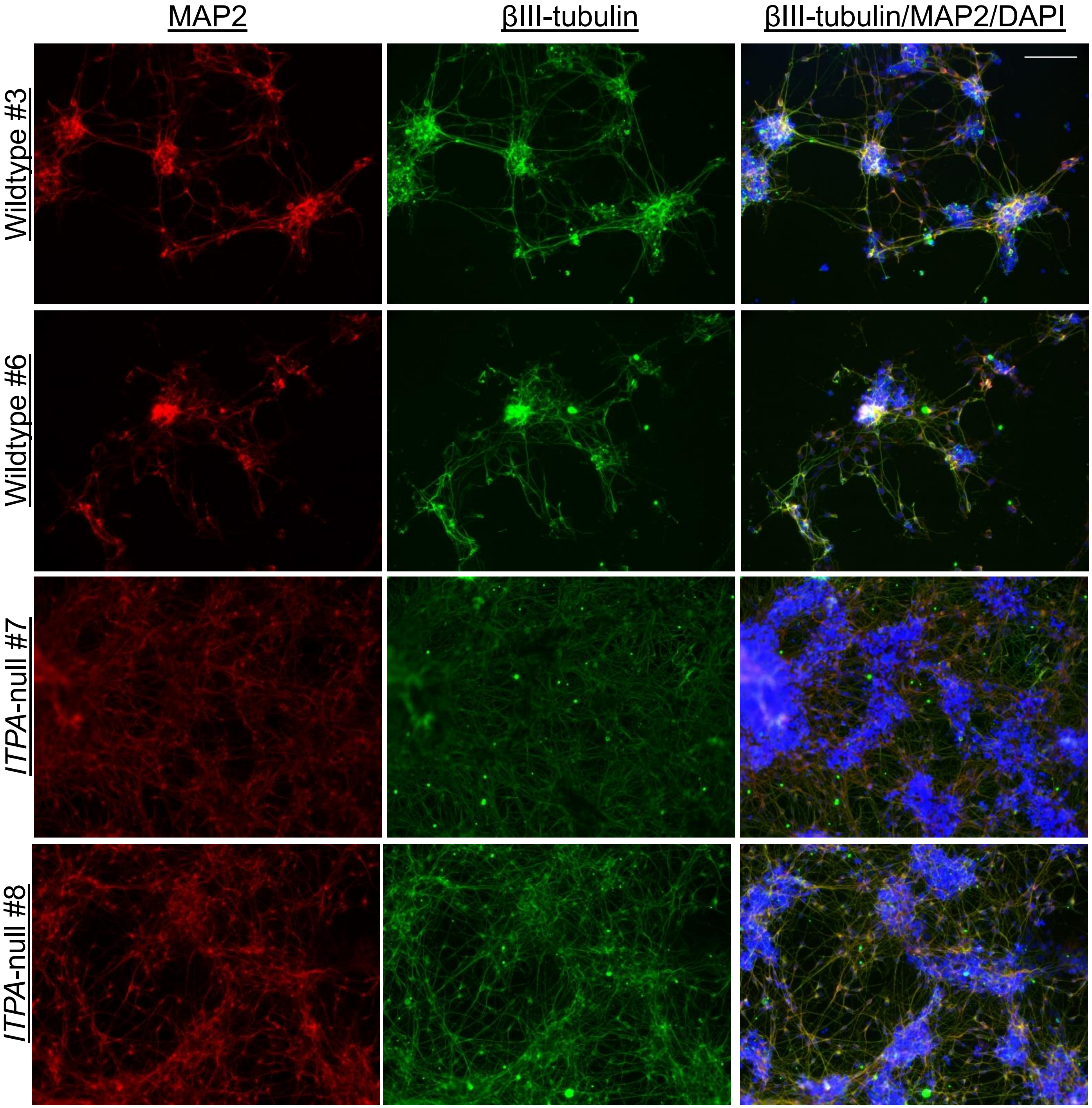


**Figure S9.** Fluorescence images of iPSC derived neurons stained for neuronal markers MAP2 and βIII-tubulin with DAPI for nuclei, scale bar = 100 μm.

**Table S3.** Primer sequences

| Transcript | Forward (5'to3') | Reverse (5'to3') | Species |
|---|---|---|---|
| FLuc (cap/pA) | GCTGCACAAAGCCATGAAG | GAAGTACTCGGCGTAGGTAATG | N/A |
| FLuc (-cap/-pA) | CGTTCGTCACATCTCATCTACC | GAGCCCATATCCTTGTCGTATC | N/A |
| RLuc | CTACGAGCACCAAGACAAGAT | CTTCTTAGCTCCCTCGACAATAG | N/A |
| IRES FLuc | CGCATGCCAGAGATCCTATTT | GCCCATATCCTTGCCTGATAC | N/A |
| SDM CUG start | GCTTGGTACCCTGAGCGAGCAGA | TTGGGTCTCCCTATAGTGAG | N/A |
| SDM AGG start | GCTTGGTACCAGGAGCGAGCAGA | TTGGGTCTCCCTATAGTGAGTCG | N/A |
| SDM UGA start | GCTTGGTACCTGAAGCGAGCAGAAG | TTGGGTCTCCCTATAG | N/A |
| *Ifih1* | GAGAACTGCTGAGAAGGATAGTG | GCCTCTGTCTCCAGACTTAATG | *R. norvegicus* |
| *Eif2ak2* | GAGAGACGAAATGGCAGAGTAG | TCCAAGAAACCGCACAGTAG | *R. norvegicus* |
| *Rig1* | GCTAACCAGATCCCTGTGTATG | GATCCTGCTCTCCTCATCTTTG | *R. norvegicus* |
| *Adar* | GGCCAGTCTCAGAAGAACTATG | CGTGCTGATGTAGAGGTGAAA | *R. norvegicus* |
| *Atf4* | CTCTCTGTACGCTGTTCCTTTC | CTCGGTCATGTTGTAGGGATTT | *R. norvegicus* |
| *Gapdh* | ATGACTCTACCCACGGCAAG | CTGGAAGATGGTGATGGGTT | *R. norvegicus* |
| *IFIH1* | TCGAATGGGTATTCCACAGACG | GTGGCGACTGTCCTCTGAA | *H. sapiens* |
| *EIF2AK2* | GCCGCTAAACTTGCATATCTTCA | TCACACGTAGTAGCAAAAGAACC | *H. sapiens* |
| *RIGI* | CTGGACCCTACCTACATCCTG | GGCATCCAAAAAGCCACGG | *H. sapiens* |
| *ADAR* | ACCAGGTGAGTTTCGAGCC | TCTTGTAGGGTGAACACCGTG | *H. sapiens* |
| *ATF4* | CCCTTCACCTTCTTACAACCTC | GGCTCATACAGATGCCACTATC | *H. sapiens* |
| *GAPDH* | AATCCCATCACCATCTTCCA | TGGACTCCACGACGTACTCA | *H. sapiens* |
